# *Adhatoda Vasica* rescues the hypoxia dependent severe asthma symptoms and mitochondrial dysfunction

**DOI:** 10.1101/2020.04.01.019430

**Authors:** Atish Gheware, Lipsa Panda, Kritika Khanna, Naveen Kumar Bhatraju, Vaibhav Jain, Shakti Sagar, Manish Kumar, Vijay Pal Singh, S Kannan, V Subramanian, Mitali Mukerji, Anurag Agrawal, Bhavana Prasher

**Author notes:** Address for reprint requests and other correspondence: Bhavana Prasher, Centre of excellence for Applied Development of Ayurveda, Prakriti and Genomics, CSIR’s Ayurgenomics Unit–TRISUTRA (Translational Research and Innovative Science ThRough Ayurgenomics), CSIR-Institute of Genomics and Integrative Biology, Delhi-110007, India. Anurag Agrawal, Center of Excellence for Translational Research in Asthma and Lung Disease, CSIR-Institute of Genomics and Integrative Biology, New Delhi 110007, India.

## Abstract

Severe asthma is a chronic airway disease that exhibits poor response to conventional asthma therapies. Growing evidence suggests that elevated hypoxia increases the severity of asthmatic inflammation among patients and in model systems. In this study, we elucidate the therapeutic effects and mechanistic basis of *Adhatoda Vasica* (AV) aqueous extract on mouse models of acute allergic as well as severe asthma subtypes at physiological, histopathological, and molecular levels. Oral administration of AV extract attenuates the increased airway resistance and inflammation in acute allergic asthmatic mice and alleviates the molecular signatures of steroid (dexamethasone) resistance like IL-17A, KC, and HIF-1α (hypoxia inducible factor-1alpha) in severe asthmatic mice. AV inhibits HIF-1α levels through restoration of expression of its negative regulator-PHD2 (prolyl hydroxylase domain-2). Alleviation of hypoxic response mediated by AV is further confirmed in the acute and severe asthma model. AV reverses cellular hypoxia-induced mitochondrial dysfunction in human bronchial epithelial cells - evident from bioenergetic profiles and morphological analysis of mitochondria. In silico docking of AV constituents reveal higher negative binding affinity for C and O-glycosides for HIF-1α, IL-6, Janus kinase 1/3, TNF-α and TGF-β-key players of hypoxia-inflammation. This study for the first time provides a molecular basis of action and effect of AV whole extract that is widely used in Ayurveda practice for diverse respiratory ailments. Further, through its effect on hypoxia-induced mitochondrial dysfunction, the study highlights its potential to treat severe steroid-resistant asthma.

**Significance Statement:** Severe asthma is a global health concern with a large fraction unresponsive to current treatment modalities involving corticosteroids. Recent findings suggest that elevated hypoxia has a critical role in severity of asthma. Here, we report therapeutic treatment with aqueous extract of *Adhatoda Vasica* (AV), an ayurvedic medicine, attenuates sever steroid insensitive asthmatic features in mice. The observed effects of AV are through inhibition of hypoxic response, both *in vivo* and *in vitro*. AV also reverses mitochondrial dysfunction, a key consequence associated with hypoxia, and asthma. This study highlights the translational potential of AV for the treatment of severe asthma and provides opportunities for its usage in other disease conditions where hypoxia is pertinent.

## Introduction

Asthma is a complex and heterogeneous disease that involves recurrent, partially reversible bronchial obstruction. Airway inflammation and airway remodelling is central to disease pathophysiology leading to airway obstruction (18, 31). With ~300 million people affected and 250,000 annual deaths, asthma significantly contributes to the global health burden (27). Initially, it was considered an allergic, eosinophilic Th2 biased disease (4). However, there exists non-Th2 asthma patho-phenotypes, where cells like Th17, Th1, and their respective cytokines play a significant role in disease pathogenesis and severity (13, 43). Corticosteroids continue to be the mainstay therapeutics for asthma (6, 8). However, non-Th2 severe asthma types poorly respond to steroids and have higher risk of exacerbations and morbidity (13, 43). Though severe asthma accounts for 10-15% of all patients, the costs in terms of healthcare management is higher than acute asthma (27).

The elevated hypoxic response has been reported in more than 90% of severe asthma exacerbations (5, 6, 8, 37). This condition is highly proinflammatory and primarily mediated through two key molecules: HIF-1α a transcription factor whose levels are elevated in hypoxia and PHD2, an oxygen sensor that degrades HIF-1α in normoxic conditions (2, 11, 24, 38). Besides, neutrophil survival and IL-17-producing CD4+ T helper cell (Th17) are induced in hypoxia or elevated HIF-1α conditions (12, 42) and are observed in severe steroid-resistant human asthmatics and mouse models (5, 9, 15, 25, 26, 29). Thus, modulation of hypoxia signalling or HIF-1α could be a promising strategy to combat hypoxia-induced severe asthmatic changes.

Plant-derived medicines have been traditionally used to prevent and treat many diseases and are a continuous source of novel drug leads. In India, *Ayurveda* is an ancient medicine system that offers a translational framework to connect physio-pathology with therapeutics. The actions of drugs are described on the basis of their effect on biological axes governed by three physiological entities called *doshas* namely *Vata*, *Pitta* and *Kapha* (33). These entities work in conjunction across systems and govern homeostasis in individuals but the same when perturbed from their threshold levels, result in disease conditions. We have shown inter-individual variability in genetic as well as expression level in *PHD2* between constitution types that differ in *Pitta* (P) and *Kapha* (K) (1, 34). Modulation of the PHD2-HIF-1α axis revealed the cellular hypoxic response to be an important modifier of asthma severity (2). We hypothesised that herbal medicines reported for use in asthma treatment and having a balancing effect of P-K perturbations might modulate the cellular hypoxic response. This effect could then be useful for reversing the severe asthma patho-phenotype.

In this study, we have tested the effect of *Adhatoda Vasica* (AV), commonly known as Malabar nut, an ayurvedic medicine indigenously used to treat various aspects of asthma. AV is from *Acanthaceae* family, a dense shrub found in all parts of India. It has a bitter and astringent taste with *Pitta-Kapha* balancing action, and described for the treatment of asthma and respiratory conditions. Vasicine and vasicinone from AV have been shown to have strong bronchodilatory and anti-inflammatory effects (3, 7, 10). In this study, we demonstrated that oral administration of aqueous extract of AV to the Ova-induced allergic mice reduces the cardinal features of asthma both at phenotypic as well as a molecular level. We also provide evidence for AV’s mechanism of action in asthma through modulation of the cellular hypoxic response. We observed that AV treatment to the asthmatic mice inhibits the increased hypoxic response by downregulating HIF-1α. Decline in HIF-1α also improved mitochondrial morphofunction. We further demonstrate that AV has therapeutic effect even in severe asthma condition that is augmented in mice by elevated hypoxic response and is non-responsive to steroids. This effect of AV against HIF-1α was also confirmed through molecular docking analysis which unravels its key components targeting the proteins of hypoxia-inflammation axes.

## Results

### AV attenuates airway pathophysiology of acute allergic lung inflammation in mouse model

We first measured the therapeutic effects of AV extract (fig. S1A and table S1 and S2) on the pathophysiological features of acute allergic airway inflammation using Ova-allergen mouse model. A schematic showing the timeline of model development and drug/vehicle treatments is shown in fig. 1A. Mice sensitized and challenged with Ova exhibited increased airway resistance, eosinophilic infiltration into the lungs, mucus metaplasia, and airway remodelling, which were reversed by dexamethasone or AV treatment (fig. 1). However, the beneficial effects of AV were found to be non-linear in the range of doses tested (fig. S1). The 130 mg/Kg (AV-D2) dose of AV was found to reverse all the aforementioned pathophysiological features, while considerable variability was observed in lower and higher doses (fig. 1, S1). For instance, Ova-induced increase in airway resistance was found to be reduced by all except the highest AV dose (fig. S1B). But histological assessment of fixed lung sections showed that AV-D2 is more effective in reducing lung inflammation (fig. 1C, F, and S1C, G), mucus metaplasia (fig. 1D, G, and S1D, H), and sub-epithelial collagen deposition (fig. 1E, H and S1E, I), compared to other doses. Assessment of eosinophils in BAL fluid and TGF-β levels in lung homogenate substantiate the efficacy of AV-D2 compared to other doses (fig. 1I, J and S1J, K). In contrast, all the AV doses considered were found to significantly reduce the levels of Th2 cytokines, IL-4, IL-5, and IL-13 (fig. 1K and Table S3) in the lung. Taken together, AV-D2 is more effective, compared to other doses, in the remission of Ova-induced pathological features.

**Fig. 1.**
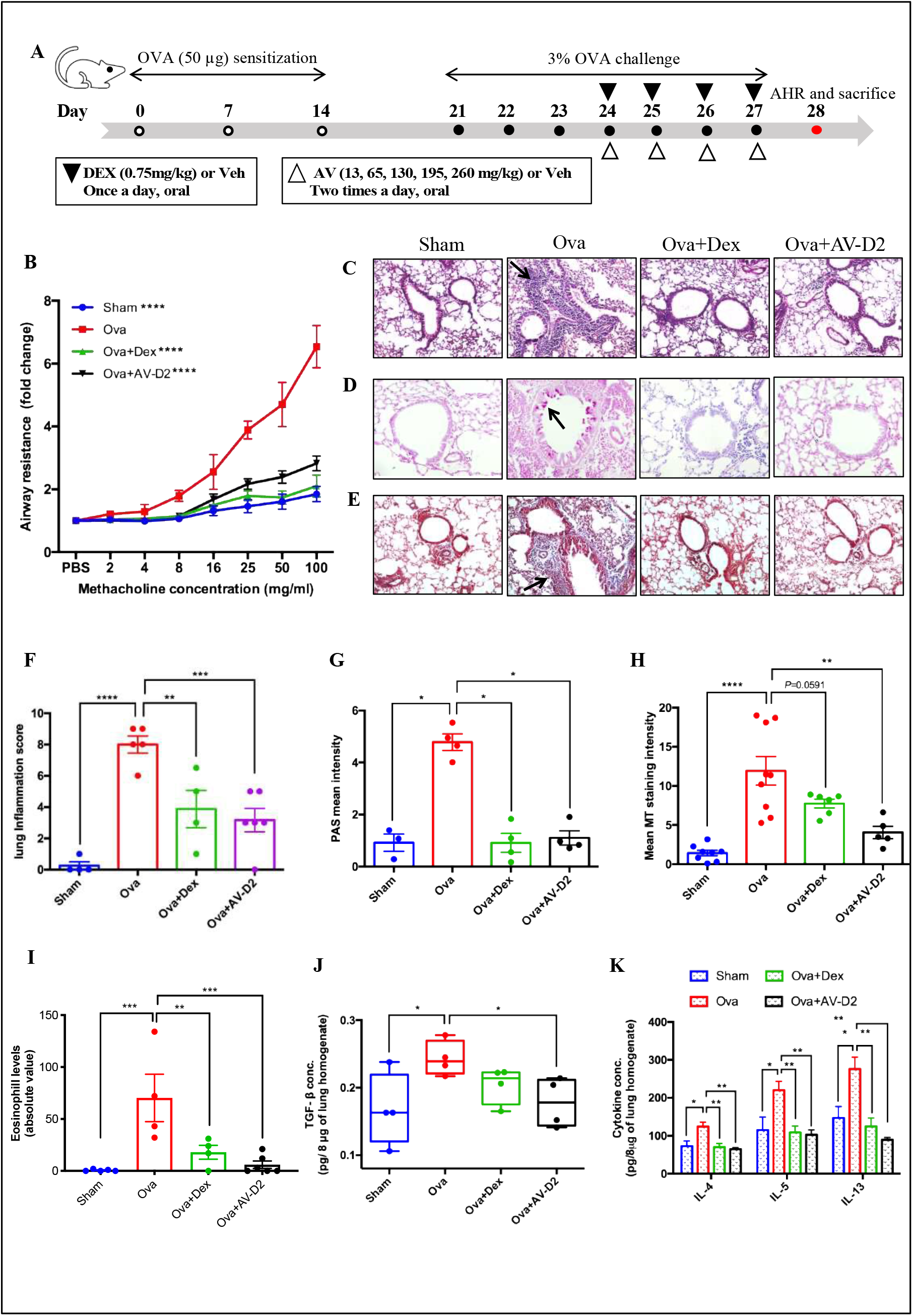
AV treatment alleviates the asthmatic features in mice model of acute asthma. **(A)** Schematic representation showing the timelines for mice model development and drug treatments. Male BALB/c mice were sensitised and challenged using Ova-allergen and AV or Dex was administered to Ova-allergic mice from day 24 to 27 as described in materials and methods. **(B)** Flexivent analysis of airway resistance after methacholine treatment in AV or Dex treated Ova allergic mice compared with Ova alone mice. **(C to E)** Representative photomicrographs of fixed mouse lung tissue sections stained with (C) H&E (10X magnification), (D) PAS (20X magnification), and (E) MT (10X magnification) for the analysis of cellular infiltration of inflammatory cell, mucin and collagen levels, respectively. Black arrow indicates positive staining in respect to particular stain. **(F)** Quantification of peribronchial and perivascular inflammation of mouse lung tissues stained with H&E in using inflammation grade scoring system. **(G and H)** Densitometric analysis of mouse lung tissues stained with PAS and MT to measure mucus metaplasia and collagen deposition using ImageJ. **(I)** Eosinophil abundance in mouse BAL fluid. **(J and K)** ELISA for TGF-β1 and Th2 cytokines in mice lung homogenate. Data are shown as mean ±SEM of three to seven mice per group and representative from three independent experiments. Significance denoted by \**P* ≤0.05, \*\**P*≤0.01, \*\*\**P*≤0.001 and \*\*\*\**P*≤0.000; by two way ANOVA (B) and ordinary one way ANOVA (F to K). **Ova-** chicken egg albumin, **Sham-**vehicle (PBS), **Dex-**Dexamethasone (0.75mg/kg), AV-**D2**-*Adhatoda Vasica* extract (130 mg/kg)

### Increased cellular hypoxia (HIF-1α) levels are reduced after AV treatment both *in vivo* and *in vitro* conditions

Our group’s earlier study has shown that the differential severity of airway inflammation in allergic asthma could be modulated by prolyl hydroxylase 2 (PHD2) inhibitor an inhibitor of HIF-1α that governs the hypoxia axis (2). Exaggerated cellular hypoxic response causes increase in disease severity in acute Ova allergic mice (2, 5). We wanted to explore the effect of AV treatment on hypoxic response elevated in Ova challenged mice. We observed elevated levels of HIF-1α in Ova mice, compared to Sham mice (fig. 2A, B, and C). Dexamethasone treatment failed to restore the increase in HIF-1α levels, whereas AV-D2 treatment completely reversed it (fig. 2A, B and C). We observed a similar non-linear effect of increasing AV doses on the HIF-1α levels as in the previous results (fig. S2A, B). In fact, a strong positive correlation (R^2^ = 0.63) was observed between airway inflammation score and HIF-1α levels with respect to D0, D2, and D4 doses of AV (fig. S2C). Further, AV treatment also restored the *PHD2* mRNA levels, which was reduced in Ova mice lungs (fig. S2D). To further validate the inhibitory effect of AV on HIF-1α levels we tested its effect on chemically induced hypoxia condition in human bronchial epithelial cells (BEAS2B). We used dimethyloxaloylglycine (DMOG), an inhibitor of PHD and FIH to induce cellular hypoxia, 10μg/ml of AV could attenuate the DMOG induced increase in HIF-1α levels in BEAS2B cells (fig. 2D, S2E). This confirms the inhibitory effect of AV on HIF-1α levels in cellular hypoxia conditions.

**Fig. 2.**
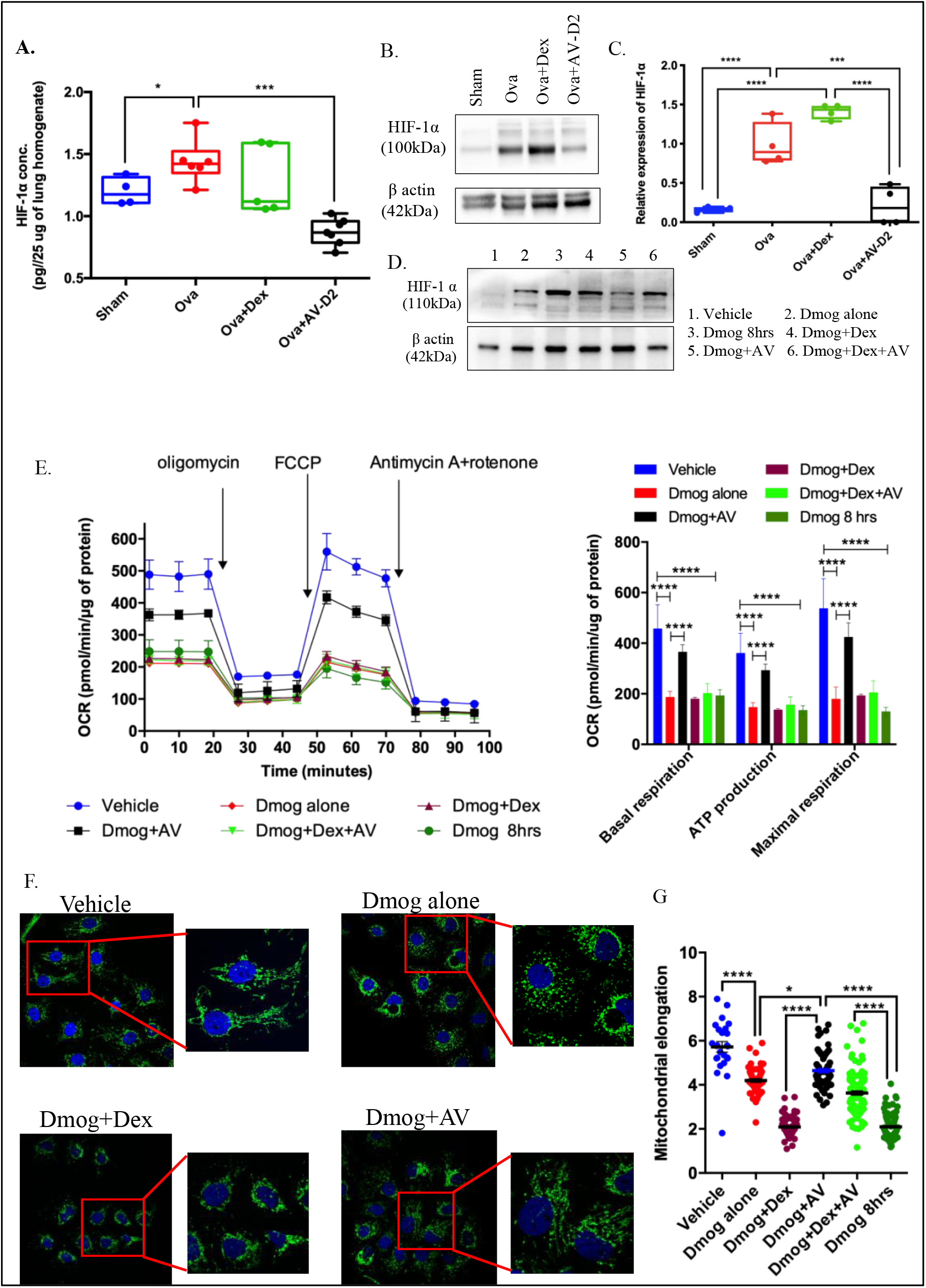
AV treatment rescues hypoxia induced HIF-1α and related mitochondrial dysfunction. **(A to C)** HIF-1α levels measured in lung homogenate of acute asthmatic mice by (A) ELISA and (B) Western blot analysis, (C) compiled with densitometric comparison for western blot. Data are shown as mean ±SEM of four to seven mice per group and representative of three independent experiments. (**D)** Representative western blot for HIF-1α abundance in BEAS2B cell lysate. (**E to G**) Effect of AV on cellular hypoxia induced mitochondrial dysfunction. (E) Left: representative OCR graph showing effect of cellular hypoxia on mitochondrial bioenergetics profile in presence of AV or Dex treatment. Right: mitochondrial bioenergetics profile measured at Basal respiration, ATP production and at Maximum respiration level in presence of ETC inhibitors. Cellular hypoxia-induced changes in mitochondrial morphology. (F) Representative confocal images showing the mitochondria in BEAS2B cells transfected with mitochondria targeted-GFP (mito-GFP) and nucleus stained with blue (DAPI). Boxed areas of image are shown with magnification to represent the morphological status of mitochondria typical to the treatment or specific condition. (G) Dot plot showing mitochondrial elongation parameter. Data are shown as mean ±SEM of thirty or more cells per group and representative of at least two independent experiments. Significance denoted by \**P* ≤0.05, \*\**P*≤0.01, \*\*\**P*≤0.001 and \*\*\*\**P*≤0.0001; by Unpaired t test with Welch’s correction (A), ordinary one way ANOVA (B), two way ANOVA with Tukey’s multiple correction test (E), ordinary one way ANOVA with Tukey’s multiple correction test (G). **Ova-** chicken egg albumin, **Sham**vehicle (PBS), **Dex**-Dexamethasone (0.75mg/kg), **AV-D2**-*Adhatoda Vasica* extract (130 mg/kg), **BEAS-2B**-normal human bronchial epithelium cells, **OCR**-oxygen consumption rate, **DMOG**-dimethyloxaloylglycine, **DMOG+AV**-DMOG+10μg/ml. *Adhatoda Vasica* extract, **DMOG+Dex**-DMOG+10nM of Dexamethasone, **DMOG+Dex+AV**-DMOG+10nM of Dexamethasone+10μg/ml. *Adhatoda Vasica* extract, **DMOG 8hrs**-DMOG treatment for 8 hours.

### Hypoxia-induced mitochondrial dysfunction is rescued by AV treatment

Hypoxia-induced mitochondrial dysfunction in asthma has been reported earlier (22, 23). Therefore, we next determined whether AV could also have an effect on mitochondrial health. Induction of cellular hypoxia in human bronchial epithelial cells (BEAS2B) by DMOG led to an overall decrease in OCR (fig. 2E) as well as mitochondrial basal respiration, ATP production, and maximum respiration. We observed similar results using adenocarcinomic human alveolar basal epithelial cells (A549). AV treatment was observed to improve mitochondrial health in both the cells, compared to vehicle group as well as in cellular hypoxic environment (fig. 2E and S3A). We next examined the effect of cellular hypoxia stress on mitochondrial morphology in BEAS2B cells transfected with mitochondria-targeted green fluorescent protein (mito-GFP). To assess the alterations in mitochondria morphology, previously reported parameters such as number of individual mitochondria, area, elongation, number of networks, and mean branch length of networks were determined. Healthy mitochondria are elongated and exhibit branched chain (network) morphology. In the control (vehicle group), we found the scores for all the aforementioned morphological parameters were higher than in the DMOG treated cells (fig. 2F, G and S2F-I). AV treatment was found to restore morphological defects induced by DMOG treatment (fig. 2F, G and S2F-I). In contrast, Dex treatment alone was found to be ineffective in restoring the DMOG induced morphological alteration in mitochondria (fig. 2F, G and S2F-I), but the combination of Dex with AV was observed to be beneficial in significantly improving majority of the morphological parameters measured (fig. 2F, G and S2F-I). This suggests AV treatment may be useful in conditions where hypoxia stress is prominent with compromised mitochondrial morphology and function. To further validate this, we tracked the mitochondria labelled with TOM20 (mitochondrial import receptor subunit) antibody in BEAS2B cells treated with DMOG or vehicle. We observed a significant improvement in mitochondrial morpho-functions (fig. S3B) after hypoxic stress. Noteworthy, this was not restored with Dex treatment (fig. S3C-F). However, the combination of Dex with AV treatment in cellular hypoxic stress could restore some of the changes in morphological aspects of mitochondria as shown in fig. S3C-F.These results establish that AV targets the hypoxia-oxo/nitrative stress-mitochondrial dysfunction axis to attenuate the airway pathology during asthma.

### Severe airway inflammation resistant to steroid is resolved with AV treatment

#### DHB induced severe asthma model

Earlier, we showed that chemical, as well as siRNA-mediated inhibition of PHD-2 in allergic mice led to increased hypoxic response and severity of the disease (2). In this study we used the same model of severe asthma to identify their steroid sensitivity (fig. 3A). We found that chemical inhibition of prolyl-hydroxylase (PHD) level in Ova-allergic mice by DHB (10mg/kg,) treatment led to severe increase in airway hyper-responsiveness (AHR) compared to mice challenged with OVA alone and this increased AHR was resistant to Dex treatment (fig. 3B). Therapeutic treatment of AV (130mg/kg, AV-D2) to such DHB+Ova mice led to a significant reduction in airway resistance compared to Ova+DHB, Ova+DHB+Dex mice group (fig. 3B). Similarly, DHB treatment also led to severe increase in airway inflammation (AI) and mucus metaplasia in mice (fig. 3C, G) which was confirmed by quantitative scoring and morphometry of mouse lung sections (fig. S4A, B). This increased AI and mucus metaplasia were resistant to Dex treatment, but AV could rescue these features (fig. 3C, G and S4A, B). DHB treatment to Ova mice also leads to a significant increase in IL-13 levels, which was significantly reduced after AV-D2 treatment and was unaffected by Dex treatment in Ova+DHB mice (fig.S4C). To confirm whether the observed steroid insensitive effect in mice was because of increase in hypoxia and/or HIF-1αlevel, we determined the effect of DHB on HIF-1α protein levels. We observed that DHB treatment to Ova-allergic mice led to significant increase in lung HIF-1αlevels compared to Ova mice (fig. 3D) which was significantly reduced with AV treatment but not by Dex treatment (fig. 3D). In addition, cellular hypoxia induced by DHB treatment leads to significant increase in steroid non-responsive and steroid resistant asthma-associated cytokines, namely IL-17A (fig.3E), KC (fig.3F, mouse homologue of IL-8), IFN-γ and TNF-α(fig. S4D, E), compared to Ova mice group. These increased cytokine levels were also significantly reduced with AV treatment whereas Dex treatment (fig. 3 D-F and S4D, E) couldn’t reduce them. There was also a significant difference in therapeutic effect accounted by AV-D2 treatment on DHB induced severe airway inflammation compared with the Dex-treated severe asthmatic mice (Ova+DHB+Dex). These results indicate that AV’s therapeutic treatment to severe allergic mice reduces airway inflammation and associated cytokine levels through modulation of HIF-1α levels.

**Fig. 3.**
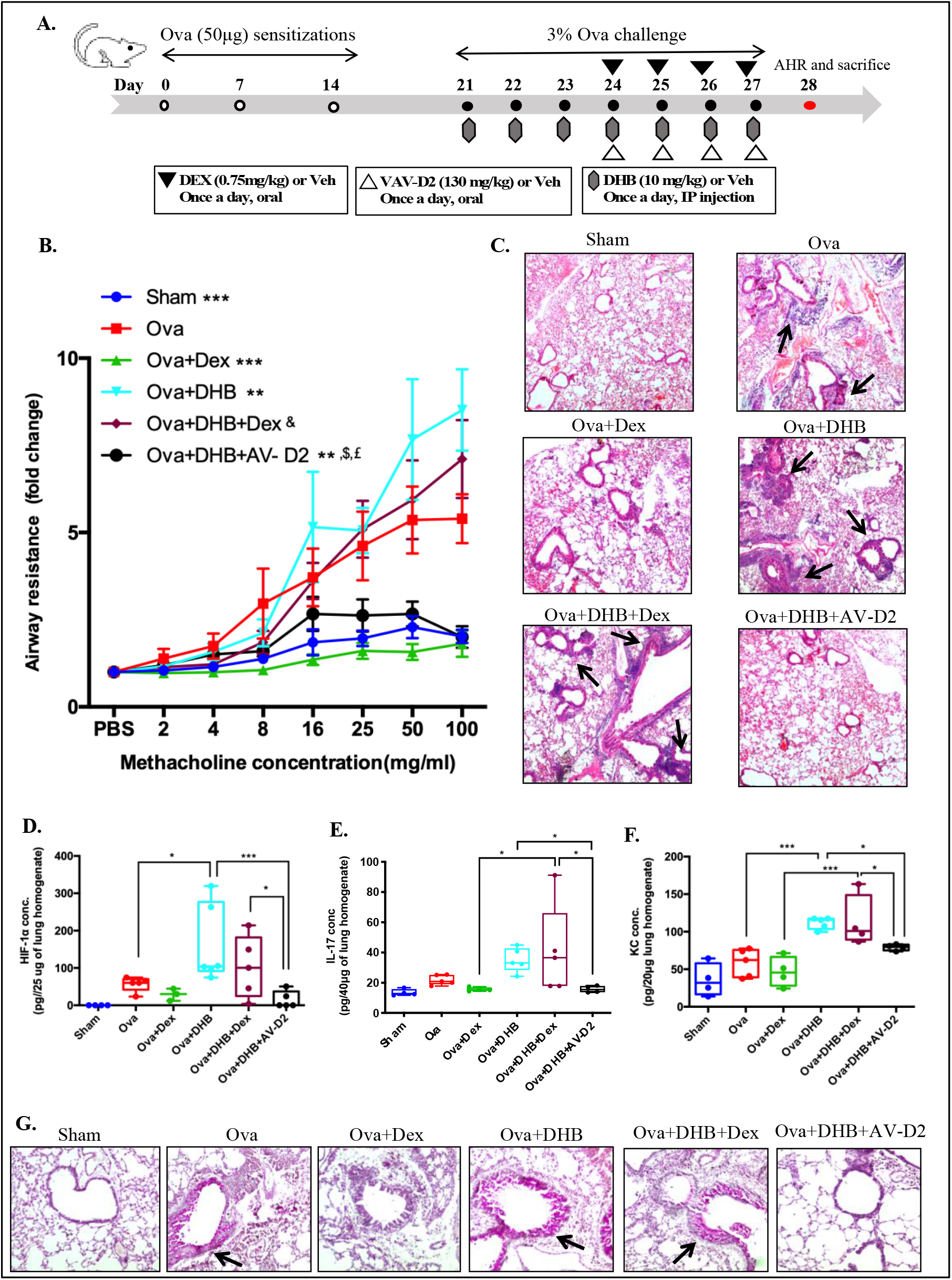
AV treatment resolves the severe steroid resistant airway pathological features induced by chemical inhibition of PHD in mice model of asthma. **(A)** Schematic representation of experimental protocol used to induce severe asthma in mice using DHB as described in materials and methods. **(B)** Flexivent analysis of airway resistance after methacholine treatment for DHB induced severe asthma mouse model. (**C)** Representative photomicrographs (4X magnification) of mouse lung tissues stained with H&E **(D to F)** ELISA analysis for HIF-1α (D), IL-17 (E) and KC (F) abundance in mouse lung tissue homogenate. **(G)** Representative photomicrographs of mouse lung tissues stained with PAS. Arrow indicates positive staining for mucin. Data are shown as mean ±SEM of three to five mice per group and representative of two independent experiment. Significance denoted by \**P*≤0.05, \*\**P*≤0.01, \*\*\**P*≤0.001 and \*\*\*\**P*≤0.0001 compared with Ova group, & denotes *P*≤0.0001 compared to Ova+Dex group, $ and £ denotes *P*≤0.0001compared to Ova+DHB group and Ova+DHB+Dex group respectively; assessed by two way ANOVA with Tukey’s multiple correction test (B), ordinary one way ANOVA (D and F), Unpaired t test with Welch’s correction (E). **Ova-** chicken egg albumin, **Sham**-vehicle (PBS), **DHB**-ethyl, 3,4 - dihydroxy benzoic acid (10mg/kg), **Dex**-Dexamethasone (0.75mg/kg), **AV-D2**-*Adhatoda Vasica* extract (130mg/kg).

#### PHD2 siRNA induced severe asthma model

To test whether AV alleviates the steroid resistance under high cellular hypoxia in the severe asthma through specific inhibition of PHD2 in lung, we administered siRNA through intranasal route in Ova allergic mice (fig. 4A). Down-regulation of PHD2 level was confirmed by measurements of its protein levels (fig. 4B). Interestingly AV treatment increases the PHD2 protein levels significantly compared to only PHD2 siRNA administered allergic mice (fig. 4B). Consequently, a significant increase in HIF-1α was also observed in PHD2 siRNA-treated Ova mice compared to scrambled (Scrm)siRNA+Ova mice group and increased HIF-1α was significantly downregulated by AV treatment but not by Dex (fig. 4C). PHD2 siRNA recapitulated the severely increased airway resistance that was also significantly reduced after AV (130mg/kg, AV-D2) treatment but not by Dex treatment (fig. 4D). PHD2 siRNA treatment to Ova mice also increased the airway inflammation (AI), mucus metaplasia, as well as IL-17A levels significantly compared to Scrm siRNA treated Ova mice (fig. 4E, F and S5A, B and F). These increased AI and IL-17A levels were significantly reduced in AV but not in Dex treatment (fig. 4E, F and S5A, B). In addition, PHD2 siRNA treatment also increases the pro-inflammatory and steroid-resistant related cytokines like IL-13, KC and IL-6 levels (fig. S5C-E). Oral administration of AV-D2 to such severe asthmatic mice reduces these elevated cytokines levels but not by Dex treatment (fig. S5C-E). This confirms that the observed severe asthmatic features in mice is because of increase in cellular hypoxic response in allergic mouse lung and it could be attenuated by AV administration. This effect could be possibly increasing lung PHD2 levels. These results indicate that AV reduces the severe steroid-resistant airway inflammation in mice with an exaggerated hypoxic response.

**Fig. 4.**
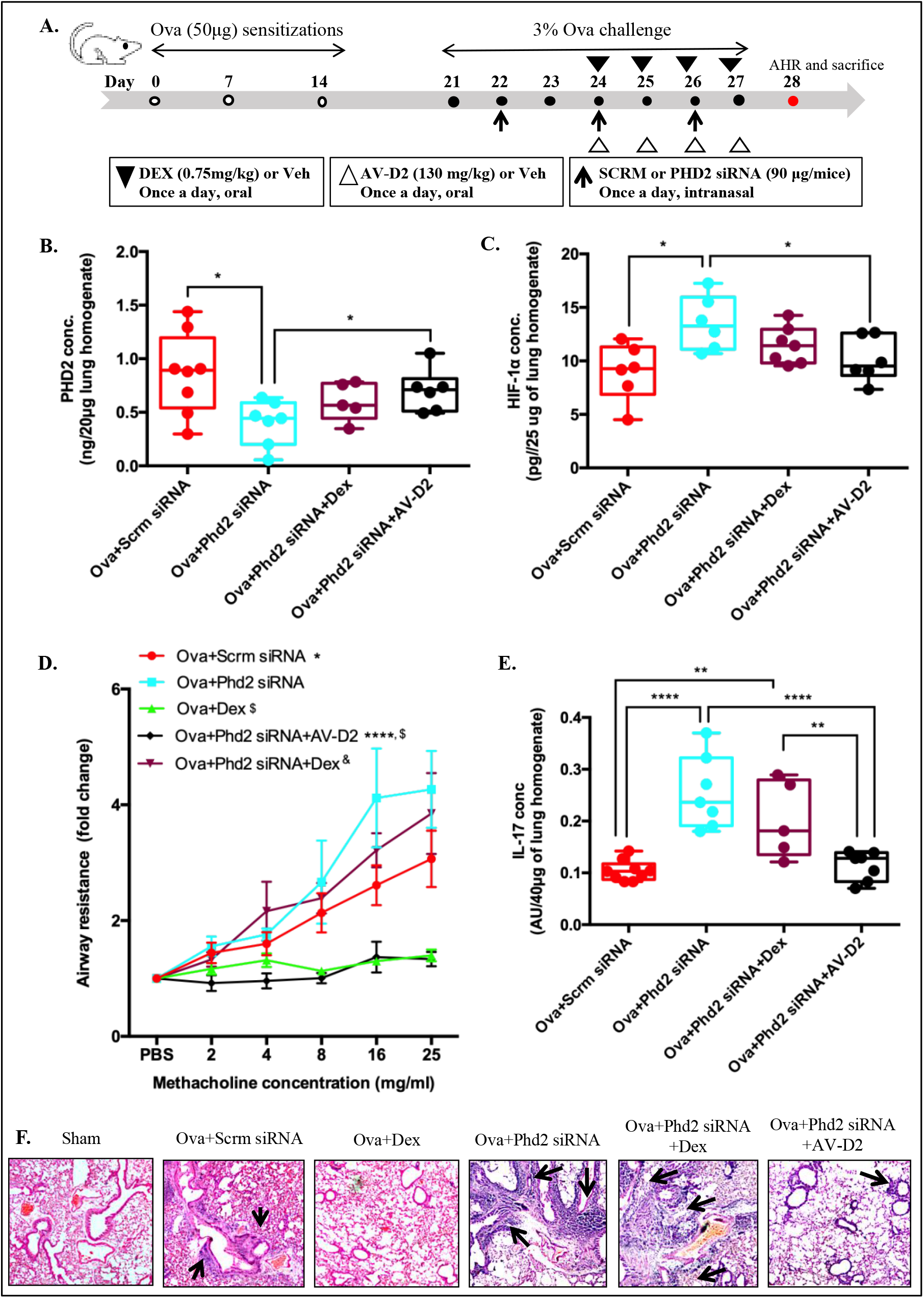
AV treatment rescues the *PHD2* siRNA induced severe steroid insensitive asthmatic features in mouse model of asthma. **A)** Schematic representation of experimental protocol used to induce severe asthma in mice using *PHD2* siRNA as described in materials and methods. **(B and C)** ELISA analysis for PHD2 (B) and HIF-1α (C) levels in mouse lung tissue homogenate. **(D)** Flexivent analysis of airway resistance after methacholine treatment in *PHD2* siRNA mediated severe asthma mouse model. **D)** ELISA analysis for IL-17 abundance in mouse lung tissue homogenate. **(E)** Representative photomicrographs (4X magnification) of mouse lung tissues stained with H&E. Black arrow indicates airway inflammation. Data are shown as mean ±SEM of five to nine mice per group and representative of two independent experiment. Significance denoted by \**P* ≤0.05, and \*\*\*\**P*≤0.0001compared with Ova+PHD2 siRNA mice, £ denotes *P* ≤0.05 compared to Ova+Scrm siRNA group, $ indicates *P*≤0.01 compared with Ova+PHD2 siRNA+Dex group and & denotes *P*≤0.0001compared with Ova+Dex group; assessed by assessed by Unpaired t test with Welch’s correction (B and C), two way ANOVA with Tukey’s multiple correction test (D), and ordinary one way ANOVA (**E**). **Ova-** chicken egg albumin, **Sham**-vehicle (PBS), **Scrm siRNA**-Scrambled siRNA-90μg/mice, **PHD2 siRNA-** mouse specific PHD2 siRNA 90μg/mice., **Dex**-Dexamethasone (0.75mg/kg), **AV-D2**-*Adhatoda Vasica* extract (130mg/kg).

### Molecular docking predicts key constituents of AV’s anti-hypoxic targets

To identify the constituent specific effect of AV extract on hypoxia-inflammation relevant targets, we performed molecular docking analysis (figure 5, S6 and table S4). The results showed luteolin-6,8-di-C-glucoside from AV extract, has a higher binding energy for HIF-1α (−9.31kcal/mol.), IL-6 (−6.94 kcal/mol.), and Janus kinase-1 (JAK-1) (−12.08 kcal/mol.). The compounds kaempferol-3-O-rutinoside and Apigenin-8-C-arabinoside of AV exhibit binding energy of −7.16 kcal/mol for IFN-γ and −11.28 kcal/mol for JAK-3 protein respectively. The luteolin-6C-glucoside-8C-arabinoside component also shows a binding energy value of −9.41 kcal/mol with TNF-α. It is interesting to note that Kaempferol-3-O-rutinoside exhibits π-π stacking interactions with the residue Trp 32 of TGF-β protein with −7.45 kcal/mol binding energy (figur S6 and table S5). The AV’s component also possess a higher binding affinity for the other proteins involved in airway inflammatory signalling cascade, such as JAK-1 and JAK-3 proteins (table S4). The key residues and number of hydrogen bonds of AV compound having higher binding free energy values are listed in table S5. In order to understand the effectiveness of whole extract of AV on severe airway inflammation due to hypoxia, the compounds of AV are docked with these proteins in other possible binding sites. As shown in figure 5, the multiple compounds of AV extract have appreciable binding affinity (numerically, more than −6 kcal/mol) in binding sites for proteins relevant to inflammation biology and cellular hypoxic response.

**Figure 5.**
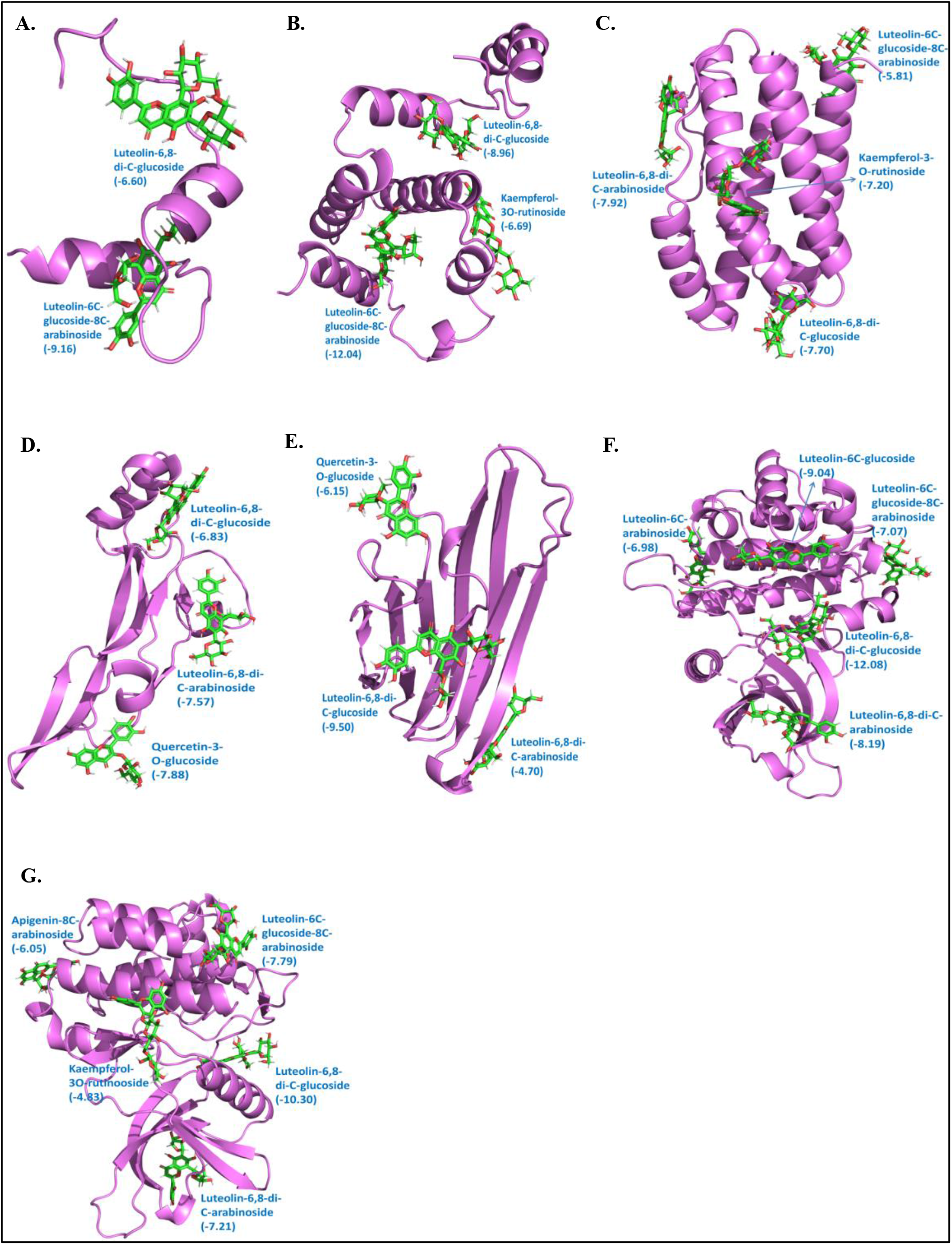
Interaction of compounds of AV with multiple binding sites of proteins related to hypoxia and inflammation pathway. (**A**) HIF-1α, (**B**) IFN-γ, (**C**) IL-6, (**D**) TGF-β, (**E**) TNF-α, (**F**) JAK-1, and (**G**) JAK-3. The binding affinity of the compounds is indicated in parentheses in kcal/mol.

## Discussion

Plant derived medicines, like phytoextracts, form an integral component of indigenous medical systems and have been successfully used to prevent and treat various diseases (17, 19, 40). In this study, we use aqueous extract of the plant *Adhatoda Vasica* (AV) which was prepared as per descriptions in *Ayurvedic* texts to understand its effect and possible mode of action against asthma (16, 30, 35). AV is one of the primary medicinal herbs in *Ayurveda* for the treatment of cough, bronchitis, asthma and various respiratory ailments (10). A number of active constituents have been isolated from AV herb, among which vasicine and vasicinone are the primary alkaloids and known to have bronchodilatory, anti-inflammatory and anti-asthmatic effects (3, 10). Semisynthetic derivatives of vasicine, bromohexine and ambroxol, are also widely used either alone or in combination with expectorant and mucolytic agents. (20). Although, none of the studies tested the detailed effect of AV on molecular markers of asthma/inflammation phenotype in its aqueous extract form as prescribed in Ayurveda. We observed, therapeutic administration of AV has a strong inhibitory action against increased airway resistance and airway inflammation observed in Ova challenged allergic mice (fig. 1). Though the therapeutic effects on the cardinal features of asthma like airway hyperresponsiveness (AHR) and airway inflammation (AI) was observed nearly at all four doses, of AV, the D2 dose was found to be most effective across all parameters with least variability within the group. Its effect on cytokine levels was dose-independent and all the doses were more or less equally effective in reducing Th2 and Th17 cytokines (table S3 and fig. S1F). However, AV demonstrated a “U-shape curve effect” observable at the level of airway physiology as evident in AHR (fig. S1C.) as well as AI (fig. 1F) and, in airway remodelling (fig. S1D, E) mediated by increased TGF-β1 in allergic mice (fig. 1H). This U shaped curve effect was mirrored in HIF-1α levels indicating the role of AV as a modulator of the cellular hypoxic response (fig. 2A).

A modifying role of elevated cellular hypoxic response in acute allergic asthma primarily mediated through hypoxia inducible factor-1α (HIF-1α) has been shown by our group earlier(2). Mild or low level of cellular hypoxia mediated by PHD2 inhibition produces the protective effect in allergic asthma whereas, high hypoxia level induces exaggerated pro-inflammatory and pro-asthmatic effects in allergic mice (2). Levels of IL-17A (a Th17 cytokine) and neutrophils are known to positively regulated by HIF-1α and both IL-17A and neutrophils are high in severe steroid resistant human asthmatics and mouse models (2, 5, 8, 9, 12, 15, 25, 29, 42). We therefore examined the effect of PHD inhibition and hypoxia response for steroid sensitivity as well the effects of AV on hypoxia induced severe asthma model. Here, we show that chemical as well as siRNA mediated induction of HIF-1α in allergic mice increases the asthma severity (fig. 3, 4). We also show that AV treatment is able to rescue all the severe asthma phenotypes including its effect on molecular markers like IL17, KC which were found to be non-responsive to the steroids (Dex) (fig. 3, 4, S4 and S5). This could be because of inhibitory effect of AV on increased HIF-1α levels or by restoration of decreased PHD2 levels in allergic severe asthmatic mice (fig. 4B, S2C), The inhibitory effect of AV on HIF-1α in cellular hypoxic condition is also validated by *in vitro* experiments (fig. 2D, S2D).

One of the axes associated with severe inflammation is hypoxia-oxo-nitrative stress-mitochondrial dysfunction in asthma (22, 23, 32, 36). Indeed, increased cellular hypoxic response has been shown to cause mitochondrial dysfunction and oxidative stress, (22, 36). We further examined if AV could improve the consequences of cellular hypoxic response such as mitochondrial dysfunction and thereby modulate the outcomes of the disease. Mitochondrial dysfunction was assessed *in vitro* in cultured lung epithelial cells treated with DMOG to induce the cellular hypoxic response. Our results reveal that AV treatment to lung epithelial cells was able to reverse the DMOG induced mitochondrial dysfunction as measured through mitochondrial morphological analysis and increased OCR (fig. 2E-G, S2E-I and S3). AV treatment restored the thread-like shape and mitochondrial network morphology altered in cellular hypoxic stress (fig. 2E-G, S2E-I and S3). The beneficial effects of AV on mitochondrial respiration could be either through direct attenuation of cellular hypoxic state or indirectly through increase in mitochondrial biogenesis (32). It would be interesting to test the effect of AV in corticosteroid insensitive obese-asthma phenotype, where the mitochondrial dysfunction and hypoxia co-exist (23).

Since AV is a multicomponent formulation, we wanted to identify the specific constitutes that could be potentially targeting the inflammatory, and hypoxia axes. Molecular docking analysis revealed that the C and O-glycosides from AV extract, such as luteolin-6,8-di-C-glucoside, have a higher negative binding affinity for HIF-1α and their downstream targets such as IL-6, TNF-α and TGF-β (Fig S1A, Table S4). This is the central axis connecting the cellular hypoxic response, via its principal mediator HIF-1α, to cellular inflammatory pathways. This pathway is also well known to be modulated by oxidative stress and therefore other than pharmacological binding, the anti-oxidant potential of such molecules may also be important. Besides, various components from AV also show a higher binding affinity for JAK1 and JAK3 proteins, essential in cytokine signal transduction to induce type 2 asthma effects (5). This observation coincides with our *in vivo* and *in vitro* results, which validate AV’s therapeutic impact on a hypoxia-inflammation axis. It is unlikely that there is a single active molecule in such herbal drugs. Thus, a deeper understanding of the mechanism of action may be explained through novel approaches like network pharmacology. (21)

Inter-individual variability in baseline levels of PHD2 may predispose individuals to severe hypoxia response in respiratory disorders. Previously, we observed genetic difference in *PHD2* between healthy individuals of *Pitta* and *Kapha* constitution types of Ayurveda between natives of high altitude and sojourners who suffered from HAPE (1). Subsequently we showed the significance of PHD2 in asthma, where chemical inhibition of PHD2 modulated hypoxic response in asthma and increases disease severity (2). The drugs prescribed in *Ayurveda* are also classified on the basis of their effect on the *Tridoshas, Vata, Pitta and Kapha* (28). The herbal medicine *Adhatoda Vasica* tested in this study is described to be *Pitta-Kapha* balancing herb and used for treatment of asthma and other respiratory conditions (28). Effect of AV on asthma not only lead to identification of novel inhibitor of HIF-1α, that could be useful in treatment of severe asthma, but also explains/validates the molecular correlate of *Pitta-Kapha* axis identified through the inter-individual variability between *Pitta-Kapha* constitution types using *Ayurgenomics* approach.

Taken together, we demonstrate for the first time that *Adhatoda Vasica* extract modulates cellular hypoxic response and has potential for ameliorating severe asthma. In addition, AV may protect against mitochondrial dysfunction in the cellular hypoxia state. As we have described, the extract (fig. S1A and table S1, S2) used in this study is multi-component in nature. Thus, it is likely that AV may act on multiple molecular targets or processes for the observed effects. While molecular docking helps in identifying some of the likely actors, further dissection may require novel approaches such as network pharmacology. Nevertheless, the results of our study highlight possible molecular mechanisms that could explain the clinical efficacy of aqueous AV extract reported in human asthmatic patients (16, 35). The anti-inflammatory effects of AV observed in acute asthma models indicate that AV could be an adjunct or alternative to steroids. This study demonstrates the potential of AV treatment specifically in severe steroid resistant asthmatics and in conditions where restoration of mitochondrial dysfunction is pertinent. The therapeutic observation of AV extract on cellular, histological (tissue level) and physiological (phenotype level) parameters involved in pathophysiology reflects the importance of retaining the multi compound nature of the extract, as practised in traditional medical systems like *Ayurveda*.

## Materials and Methods

AV was prepared according to the classical method described for *rasakriya* (decoction-condensation-drying) in *Caraka Samhita* (35, 39). The detailed quality and chemical fingerprinting study were carried out by LC-MS at CSIR-CDRI, Lucknow, India (fig. S1A, B and table S1, S2). Acute allergic asthma was developed in mice using Ova sensitization by intraperitoneal injection from a 1-3 week. After a 1week gap mice were challenged with Ova using an aerosol nebulizer for 7 days. AV (13-260mg/kg, orally), and Dex (0.75mg/kg, orally) was given for 4 days from the 24th-day protocol. On the 28th-day, mice were cannulated to determine airway resistance in response to methacholine and sacrificed by phenobarbital anesthesia. Severe asthma model was developed by intraperitoneal injection of DHB (10 mg/kg), and intranasal PHD2 siRNA (90 μg) treatment to induce a hypoxic response in Ova-sensitized and challenged mice. Lungs were removed, fixed in formalin, embedded in paraffin, and sectioned at 5μm. Staining of hematoxylin and eosin, periodic acid-Schiff, and Masson Trichrome was used to assess the lung inflammation, mucus hypersecretion, and sub-epithelial fibrosis, respectively. Quantitative real-time reverse transcription PCR (RT-PCR) was used to quantify PHD2 mRNA expression. Lung tissue lysate was used for cytokines and western blot assay. Cellular hypoxia model was developed using DMOG (1mM; 32 hours), in BEAS2B and A549 lung epithelial cells. AV(10μg/ml), and Dex (10μM/ml) were added to culture after 8 hours of DMOG induction and kept for next 24 hours. The cell lysate was used to detect HIF-1α protein. Seahorse assay was carried out using a 24-well plate with BEAS-2B, and A549 cell culture in presence of DMOG. GFP-mito or immunofluorescence was used to detect mitochondria and its morphological characteristics. Molecular docking of AV constituents with HIF-1α, IFN-γ, IL-6, TNF-α, TGF-β, JAK-1 and JAK-3 proteins was performed using Schrodinger suite (Maestro) and AutoDock vina package (14, 41). The trend from the docking results obtained from Schrodinger suite of packages is almost similar to those of AutoDock vina package. For brevity and further analysis, results from Schrodinger package are only considered. Data were analyzed by one-or two-way analysis of variance (ANOVA), Student’s t-test, and correlation analysis. Mitochondrial quantitative analysis was performed by using image J. The difference was considered to be statistically significant when *P*< 0.05.

Details of methods and experimental protocol is provided in supplementary materials and method

## Acknowledgments

We acknowledge animal house and imaging department for access to facility. We thank Dr. Ramniwas Prasher for his valuable suggestions regarding AV extract preparation and its use in clinical terms of *Ayurveda*. We also thank Dr. Balaram Ghosh, Dr. Ullagnath Mabalirajan, Dr. Krishnendu Chakroborty and Dr. Soumya Sinha Roy for discussion and suggestions in study. AG, LP and KK acknowledge AcSIR (Academy of Scientific and Innovative Research) for PhD. registrations and CSIR (Council of Scientific and Industrial Research) for fellowship.

## Funding

This work is supported by grant to CSIR-TRISUTRA (MLP-901) from the CSIR and Center of Excellence grant by Ministry of AYUSH, Govt. of India.

## Author Contributions

A.G. designed and performed the experiment, analysed the results, and wrote the paper. A.G., L.P., and K.K. performed the experiment, analysed the results. V.J. and V.P.S. contributed to animal model experiments. N.K.B. contributed to the seahorse experiment and quantification of flurosense labeled mitochondrial images. S.S. & M.K. contributed to an imaging experiment. S.K. and V.S. performed molecular docking and their analysis and interpretation. B.P. conceptualized the study, provided A.V., quality control information, discussion, designed the experiment, analysed the results, and wrote the paper. M.M. and A.A. designed experiments, analysed, and discussed the results, and wrote the paper. All the authors reviewed and approved the final version of the manuscript.

## Notes

### Competing Interest Statement

The authors have declared no competing interest.

### Summary of Updates

1) Manuscript updated with new results, where constituents specific effect of Adhatoda Vasica is explored on hypoxia-inflammation axes using molecular docking analysis. 2) Revised version updated with supplementary data.

