## supplementary information for "*Adhatoda Vasica* rescues the hypoxia dependent severe asthma symptoms and mitochondrial dysfunction"

Supplementary Information Text

**Materials and Methods**

**Preparation of plant extract and LC-MS fingerprinting:** *Adhatoda Vasica* (AV) was collected from Delhi-NCR region, India in the flowering season (November to March). Water extract of plant (leaves, twigs and flowers) was prepared according to classical method described for rasakriya in *Caraka Samhita* (6). The process for the formulation involved preparation of decoction condensation and drying as described in earlier study (8). Chemical fingerprinting of prepared AV extract was carried out by LC-MS at CSIR-CDRI, Lucknow, India; in two independent experiment. Briefly, Liquid chromatography-electrospray ionization-mass spectrometry of AV extract was recorded in positive- and negative- ion modes using an Agilent 6520 QTOF-MS/MS system coupled with an Agilent 1200 HPLC (Agilent technologies, USA) via an ESI interface **(Table S1 and S2)**. HPLC separation was carried out on a Supelco Discovery HS C18 column (15 cm×4.6 mm, 3μm) operated at 25°C. The mobile phase, which consisted of a 0.1% formic acid aqueous solution and acetonitrile. The analyses were performed on an Agilent 1200 HPLC system consisted of a quaternary pump (G1311A), online vacuum degasser (G1322A), auto sampler (G1329A), thermostatted column compartment (G1316C) and diode-array detector (G1315D). In the ESI source, nitrogen was used as drying and collision gas in both positive and negative ion mode. Detection was carried out within a mass range of m/z 50-2000 and resolving power above 15000 (FWHM). The chromatographic and mass spectrometric analyses were performed by using Mass Hunter software version B.04.00 build 4.0.479.0 (Agilent Technology, USA).

**Animals**

All animals were maintained as per the guidelines of the Committee for the Purpose of Control and Supervision of Experiments on Animals (CPCSEA). The BALB/c male mice (8-10 weeks old) were obtained from Central Drug Research Institute, Lucknow, India and were acclimatized to animal house environment one week prior to starting the experiments at CSIR-Institute of Genomics and Integrative Biology (IGIB), Delhi, India; as per the protocols approved by Institutional Animal Ethics Committee of CSIR-IGIB, Delhi, India. All the surgical procedures were performed under sodium pentobarbital anesthesia and maximum efforts were taken for minimum suffering of animals.

**Grouping of mice**

Mice were divided into two groups, acute and severe as per allergen challenge and treatment protocol. In both groups, there were three main sub-groups (*n* = 4-7): Sham (mice that were PBS sensitized, PBS challenged), Ova (mice that were Ova [grade V chicken egg Ovalbumin, Sigma] sensitized, Ova challenged and treated with vehicle, 50 % ethanol and distill water), Ova+Dex (allergic mice treated with Dexamethasone [0.75mg/kg], dissolved in 50% ethanol, by orally). In acute model, in addition to the above groups, mice were divided into five different Ova+AV groups (allergic mice treated with *Adhatoda* extract [AV, 13mg/kg, 65mg/kg, 130mg/kg, 195mg/kg and 260mg/kg], dissolved in distil water, oral). For severe model, Ova sensitized and challenged mice were further sub-divided according to treatment: Ova+DHB (allergic mice treated with ethyl 3,4-dihydroxybenzoic acid [DHB, 10 mg/kg], dissolved in 50% ethanol, intraperitoneal injection) or Ova+ scrambled siRNA (mice that were Ova sensitized, Ova challenged and administered with intranasal scrambled siRNA[90 μg]) and Ova+PHD2 siRNA (allergic mice treated with intranasal prolyl hydroxylase domain 2 siRNA and treated with vehicle), Ova+DHB+Dex (allergic mice treated with DHB [10 mg/kg] and administered with Dexamethasone [0.75mg/kg]) or Ova+PHD2+Dex (allergic mice treated with intranasal PHD2 siRNA [90 μg] and administered with Dexamethasone [0.75mg/kg]), Ova+DHB+AV (allergic mice treated with DHB [10 mg/kg] and administered with AV [130 mg/kg] or Ova+PHD2+AV(allergic mice treated with intranasal PHD2 siRNA [90 μg] and administered with AV [130 mg/kg]).

**Sensitization, Challenge, and Treatment of Mice**

In all models, mice were sensitized on days 0, 7, and 14 with 50 mg Ova (Sigma, Missouri, USA) adsorbed in 4 mg alum or 4 mg alum alone and were challenged from day 21 to 27 with 3% Ova in PBS or PBS alone consecutively, as described earlier. Acute Ova model effect was determined by three independent experiments and both severe model effect was determined by two independent experiments. Dexamethasone (sigma) was dissolved in 50% ethanol and was administered orally (0.75mg/kg) to mice from fourth day of challenge (24th day) till the last day of challenge (27^th^ day), once a day. Similarly, AV (dissolved in distilled water, was given orally (13mg/kg, 65mg/kg, 130mg/kg, 195mg/kg and 260mg/kg and denoted hereafter as AV-D0, AV-D1, AV-D2, AV-D3 and AV-D4, respectively) by gavage from day 24 to 27, once a day. DHB (TCI, Tamilnadu, India) was administered from day 21 to 27 by intraperitoneal injection (10 mg/kg) in 200µl volume of 50% ethanol, was given 2 hours before the Ova challenge. For PHD2 siRNA model, scrambled (Sigma) or PHD2 siRNA (Sigma) was dissolved in ultrapure DNAse and RNAse free water with in vivo-jetPEI as the transfection reagent (Polyplus), was administered intranasally in 90μg concentration to isoflurane-anesthetized mice 2 hours prior to Ova challenge, on day 23, 25 and 27.

**Measurement of airway hyperresponsiveness, bronchoalveolar lavage fluid collection, sera separation and histopathology**

Airway hyperresponsiveness (AHR) in response to methacholine (Mch, Sigma) was determined in pentobarbital anesthetized mice using flexivent system (Scireq, Canada), as described previously. The results were expressed in the fold change of airway resistance with increasing concentrations of Mch, considering the PBS aerosol induced airway resistance as baseline values. After the AHR measurement, bronchoalveolar lavage fluid (BAL) was collected by instilling 1 ml PBS into the tracheotomised airway and recovered BAL fluids were processed to get cell pellet that will be stained with Leishman stain to determine differential cell count as well as total cell count, as described previously. Blood was withdrawn by cardiac puncture, and serum was separated by centrifugation at 1500 ×g for 10 min and was kept at −70 °C till the measurements of IgE. Lungs were removed and fixed with 10% formalin (1, 2). Fixed lungs were further processed and embedded with paraffin. 5-mm paraffin embedded lung sections then stained with haematoxylin and eosin, periodic acid-Schiff, and Masson Trichrome staining to assess the lung inflammation, mucus hypersecretion and sub-epithelial fibrosis, respectively. Microphotographs were taken by Nikon microscope with camera (Model YS-100). The inflammation scoring was performed as per inflammation grade system by experimentally blind experts to find out the perivascular (PV), peribronchial (PB) lung inflammation to calculate the lung inflammation score (1, 5, 7).

**Measurement of Interleukin IL-4, IL-5, IL-13, IL-17, TGF-β1, IFN-γ, PHD2 and HIF-1α**

Lung tissue homogenates were used for sandwich ELISA. IL-4, IL-5, TGF-β1, IFN-γ (BD Biosciences), IL-13 (R &D), Il-17 (ebioscience), PHD2 and HIF-1α (USCN), were measured as per manufacturer’s instructions and results were expressed in picograms.

**Western Blotting**

For western blot lung tissue lysate was used. Proteins was separated on 8-10 % sodium dodecyl sulphate-polyacrylamide gel electrophoresis (SDS-PAGE), transferred onto polyvinylidene fluoride (PVDF) membranes (Millipore Corp, USA). Transferred membrane were blocked with blocking buffer (5% or 10% bovine serum albumin “BSA” in phosphate buffered saline with tween 20) or with non-animal protein blocker (NAP, G biosciences). Incubated with primary antibody (abcam, USA or ebiotech, USA) in 1:1000 dilution, followed by HRP conjugated secondary antibody and detected with DAB-H_2_O_2_ (Sigma, USA) or by chemiluminescence (ECL) method. α-tubulin (Sigma, St. Louis, MO, USA) or β-actin was used as a loading control. Signals were detected by spot densitometry (Image J software).

**Transfection, Immunofluorescence and imaging**

Human bronchial epithelial cells (BEAS2B) were seeded on glass bottomed dish in 6-well plate and cultured in bronchial epithelial cell growth medium. Cells were transfected with GFP-Mito(mito-GFP) was purchased from addagene as per manufactures instruction. This was generous gift from Dr. Shital (CSIR-IGIB, Delhi). Similarly for immunofluorescence, after appropriate induction, cells were fixed with 2% paraformaldehyde and permeabilized with 0.1% Triton-X100 (Sigma). Blocking was done with 1.5% BSA. These cells then were labeled with TOM 20 primary antibody (Santacruz), followed by Alexa Fluor 488 (Invitrogen) conjugated secondary antibodies. DAPI (Invitrogen) was used for nuclear staining.

The fluorescent images were collected using point scanning laser confocal microscope system (Nikon A1R-HD, Japan; Leica TCS SP8, Germany) with 60X, 1.4N oil immersion objective lens. The imaging software NIS element AR 5.11.01 and LAX3.1.5 was used to process the raw recorded image data. Modified version of the mitochondrial morphology plugin of ImageJ was used for automated mitochondrial morphometric analysis (4, 10, 11).

***In Vitro* Culture of Human Bronchial and Alveolar Epithelial cells**

Human bronchial epithelial cells (BEAS 2B) and adenocarcinomic human alveolar basal epithelial cell line (A549) was obtained from ATCC and cultured in DMEM high glucose medium (obtained from Sigma). For the induction of cellular hypoxia stress cells were treated with DMOG (1 mM/ml, Dimethyloxallyl Glycine, Cayman, 71210) or vehicle (DMSO and distill water) as control. After 8 hours of DMOG treatment, cells were treated with AV extract (10μg/ml), and Dex (10μM/ml). After 24 hrs o treatment, cells were harvested and the levels of HIF-1α was determined in cell lysates (USCN life Science, China) by western blot.

**Seahorse assay experiment**

Cells were seeded in a 24-well plate of the seahorse format (sup fig), with a seeding density 40000 and 60000 cells per well per 100 μl media for BEAS-2B and A549 respectively. The wells A1, B4, C3 and D6 served as blank and were left without any cells; only media was added to these wells. After 5 hours, when the cells adhered and almost attained morphology, additional 900μl media was added to each well. For the seahorse experiment, following groups were set up: Vehicle (DMSO and distil water), DMOG alone , a DMOG (1mM; 8 hours), AV extract (24 hours), DMOG+Dex (10μM/ml of Dex was added to cells after 8 hours of DMOG treatment and incubated for further 24 hours), DMOG + AV extract (AV extract is added to cells after 8 hours of DMOG treatment and incubated for further 24 hours), and DMOG+Dex+AV. The next day, in fresh media, 1mM DMOG and corresponding volume of DMSO was added to the designated wells. After 8 hours, AV extract (10μg/ml) was added to the respective wells. After 24 hours of AV extract induction, seahorse experiment was initiated to test mitochondrial function and it was determined by XF Cell Mito Stress Test assay using manufacturer’s instructions. A day prior to the seahorse experiment, a sensor cartridge was hydrated in Seahorse XF Calibrant at 37°C in a non-CO_2_ incubator overnight. On the day of the experiment, assay medium was prepared by adding 1mM pyruvate, 2mM glutamine and 10mM glucose to the basal medium. The pH of the medium was adjusted using 0.1N NaOH and warmed to 37°C until use. After the 24 hours incubation of cells with AV extract, the cell culture media was replaced with the prepared assay medium for 1 hour. After this, the wells were washed thrice with the assay medium. Meanwhile, antibiotics were prepared and added to respective ports in a constant volume mode. Following were the final concentrations at which the inhibitors was used: Oligomycin 1μM, FCCP 2μM, Rotenone 0.5 μM. Once the plate containing cells is ready and the ports have been loaded with the inhibitors, it was placed in the seahorse instrument and run. After the run, the data was normalised with protein content in each well and analysed further on.

**Molecular Docking**

The available 3D crystal structures of hypoxia and inflammation pathway proteins such as hypoxia-inducible factor-1 alpha (HIF-1α) (1L8C), transforming growth factor-beta (TGF-β) (2TGI), interferon gamma (IFN-γ) (1FG9), janus kinase-1 (JAK-)1 (6BBU), Janus kinase-3 (JAK3) (3LXK), interleukin-6 (IL-6) (1ALU) and tumour Necrosis Factor alpha (TNF-α) were taken from protein data bank (PDB) (3). The active regions of the proteins were identified by COACH meta-server and the results were compared with results from CASTp web server (9). In Schrodinger suite, all the target proteins were prepared using protein preparation wizard that included optimization followed by minimization of heavy atoms of proteins. The energy minimized 3D structures of all the ligands were prepared using LigPrep. The best pose of ligands that fit well in the protein cavity was carried out using OPLS3 force field with Glide package in Extra Precision mode (XP) mode. According to the size of binding cavity of the proteins, the coordinates x, y and z of the grid box were chosen with the grid resolution of 1 Å for calculations using AutoDock vina package.

**Statistical analysis**

All data represents mean ± SEM; n= 3-7 each group; *p <0.05, **p <0.01, ***p <0.001. p-value > 0.05 is considered non-significant (NS). Statistical significance of the differences between paired groups was determined with a two-tailed Student's t test. One-way or two way analysis of variance was used to compare multiple groups and was evaluated further with a nonparametric Mann-Whitney rank-sum test or Kruskal - wallis test wherever appropriate.

**Supplementary figures**

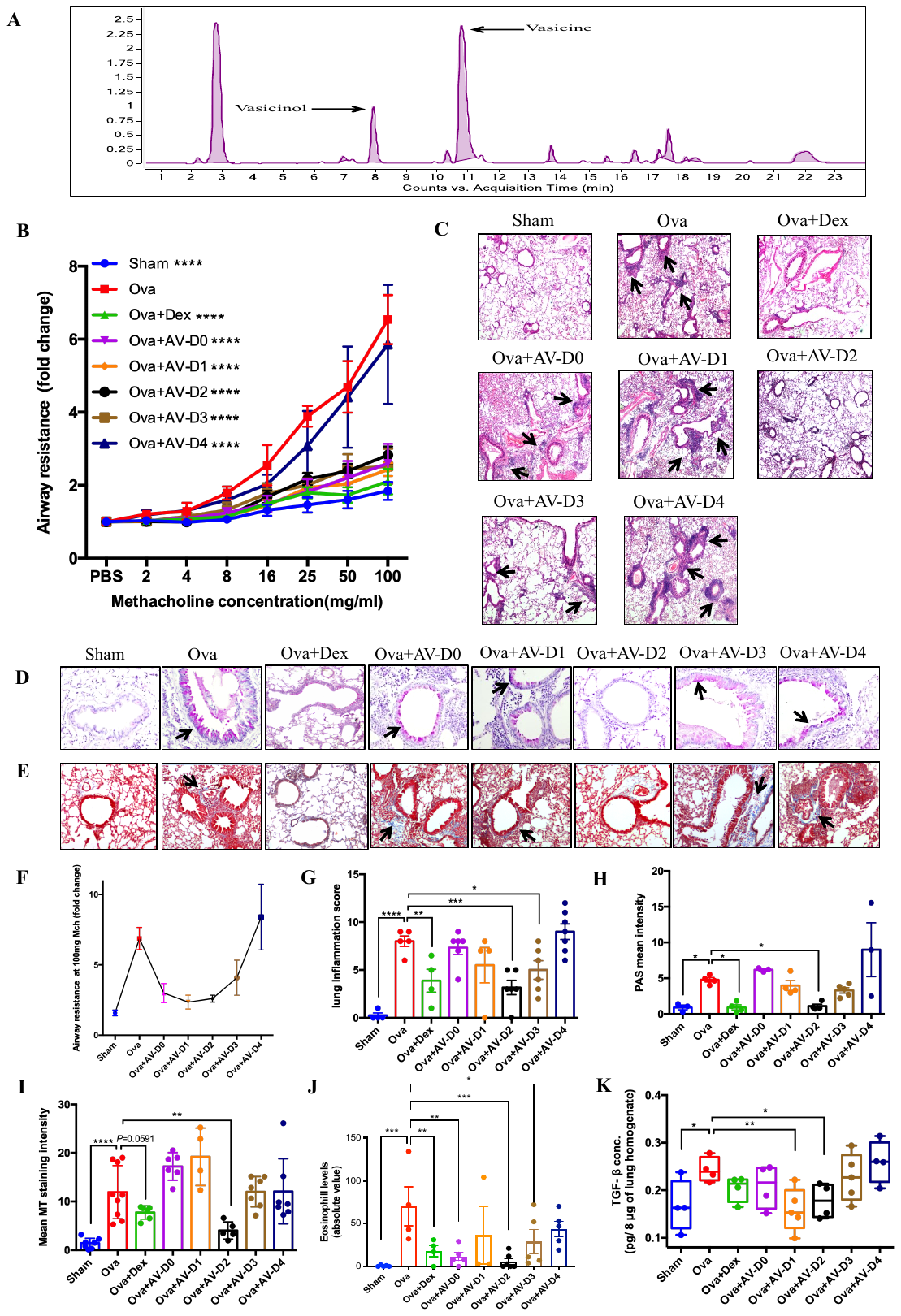

**Fig. S1. Non-linear effect (U shape curve effect) of AV on physiological parameters of acute asthma**. **(A)** LC-MS analysis of aqueous extract of *Adhatoda Vasica* using (+)-ESI-MS for identification of Quinazoline alkaloids. **(B and F)** AV shows U shape curve effect on increased airway resistance in Ova allergic mice in dose dependent manner from D0 to D4. **(C to E)** Representative photomicrographs of mouse lung tissues stained with (C) H&E (4X magnification), (D) PAS (10X magnification), and (E) MT (10X magnification) staining respectively. Black arrow indicates positive staining in respect to particular stain. (**G)** Quantification of peribronchial and perivascular inflammation of lung tissues stained with H&E in using inflammation grade scoring system. **(H and I)** Densitometric analysis of mouse lung tissues stained with PAS and MT to measure mucus metaplasia and collagen deposition respectively using ImageJ. (**J)** Eosinophil abundance in mouse BAL fluid. (**K)** ELISA analysis for TGF-β1 levels in mice lung homogenate. Data are shown as mean ±SEM of three to seven mice per group and representative from at least two independent experiments. Significance denoted by **P* ≤0.05, ***P*≤0.01, ****P*≤0.001 and *****P*≤0.000; by two way ANOVA (B) and ordinary one way ANOVA (G to K). **Ova-** chicken egg albumin, **Sham**- vehicle (PBS), **Dex**- Dexamethasone (0.75mg/kg), **AV-***Adhatoda Vasica* extract, **D0**- AV 0.130mg/kg, **D1**- AV 65mg/kg, **D2**- AV 130 mg/kg, **D3**- AV 195 mg/kg and **D4**- AV 260mg/kg.

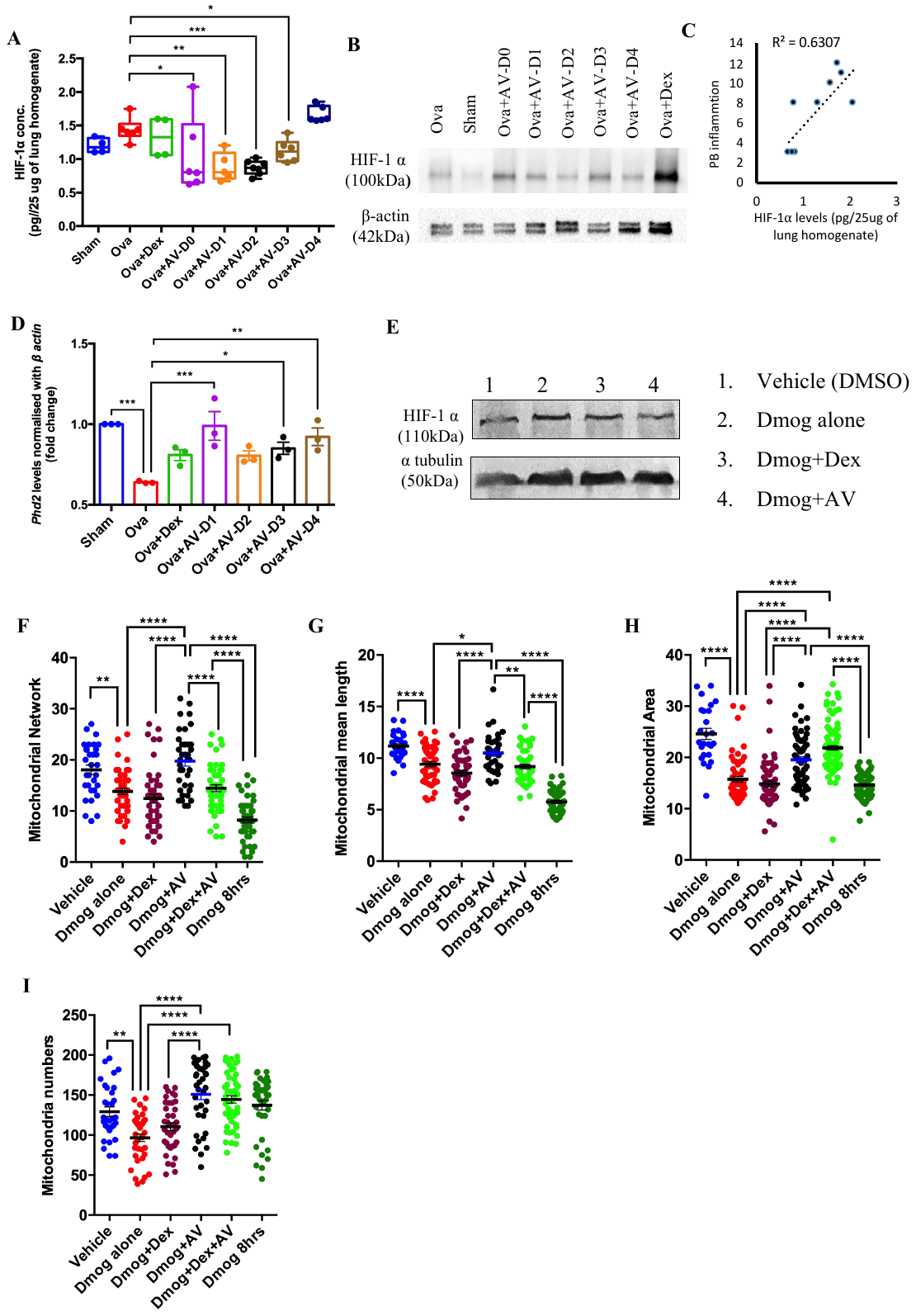

**Figure S2: AV restores the HIF-1⍺ induced increased airway inflammation in mitochondria dependent manner**. (A) ELISA analysis for HIF-1⍺ levels in mice lung tissue homogenate (*n* = 3 to 7). **(B)** Representative western blot for HIF-1⍺ abundance in lung tissue lysate of Ova allergic mice treated with Dex or AV in dose dependent manner. **(C)** Correlation analysis of airway peribronchial inflammation with HIF-1⍺ levels in mice lung after AV-D0, D2, and D4 treatment. **(D)** qPCR for *PHD2* mRNA levels in mice lung RNA samples. Data are shown as mean ±SEM (A to D). **(E)** Western blot for HIF-1α abundance in BEAS2B cell lysate. **(F to I)** Morphological analysis of mitochondria of BEAS2B cells transfected with mitochondria targeted-GFP (mito-GFP) for assessment of mitochondrial network (F), mean branch length (G), area (H), its individual number (I). Data are shown as mean ±SEM of thirty or more cells per group and representative of two independent experiments (E to I). Significance denoted by **P* ≤0.05, ***P*≤0.01, and ****P*≤0.001; by ordinary one way ANOVA (A), Unpaired t test with Welch’s correction (D), ordinary one way ANOVA with Tukey’s multiple correction test (F to I). **Sham**- vehicle (PBS), **Dex**- Dexamethasone (0.75mg/kg), **AV-***Adhatoda Vasica* extract, **AV-D0**- AV 0.130mg/kg, **AV-D1**- AV 65mg/kg, **AV-D2**- AV 130 mg/kg, **AV-D3**- AV 195 mg/kg and **AV-D4**- AV 260mg/kg, **Veh-** Vehicle (sterile distill water +DMSO), **BEAS-2B**- normal human bronchial epithelium cells, **DMOG**- dimethyloxaloylglycine, **DMOG+AV**- DMOG+10µg/ml. of AV, **DMOG+Dex**- DMOG+10nM of Dex, **DMOG+Dex+AV**-DMOG+10nM of Dex+10 µg/ml of AV**, DMOG 8hrs**- DMOG treatment for 8 hours.

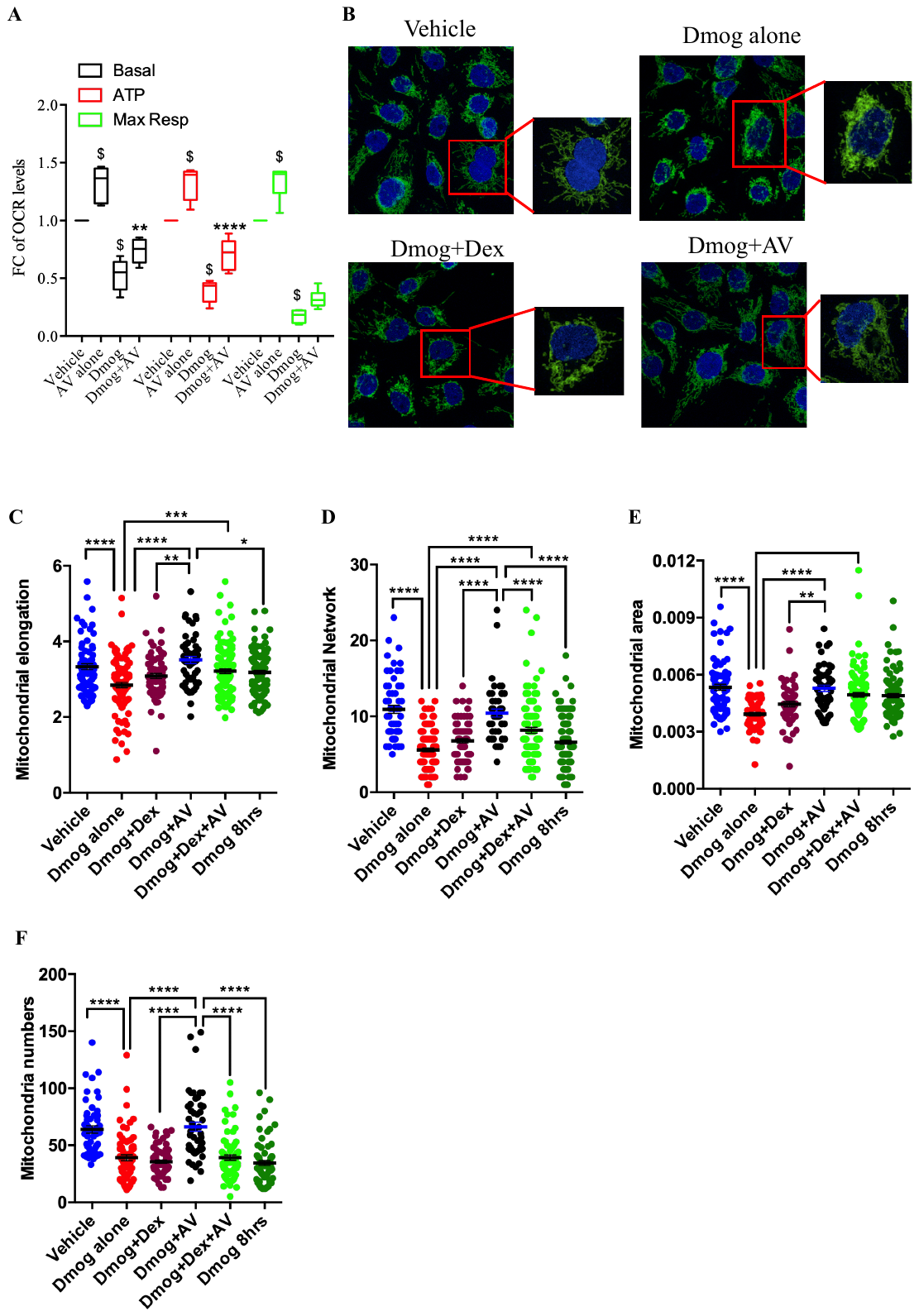

**Figure S3: AV treatment rescues cellular hypoxia induced mitochondrial dysfunction in adenocarcinomic human alveolar basal epithelial cells. (A)** Representative OCR levels linked with Basal respiration, ATP production and at Maximum respiration indicated by fold change value as compare to vehicle (sterile distil water) group in adenocarcinomic human alveolar basal epithelial cells (A549). Significance denoted by $ indicates *P*≤0.0001 compared to Veh group and ***P*≤0.01 and *****P*≤0.0001 compared DMOG group. **(B)** Representative confocal images of cells labelled with mitochondria specific TOM20 show in green, with nuclear stain (DAPI) in blue. Boxed areas in the image are magnified to show typical changes in mitochondria after each treatment and condition. **(C to F)** Dot plot showing statistical score of mitochondrial elongation (C), mitochondrial network (D), area (E) and its individual numbers per cell (F). Data are shown as mean ±SEM of thirty or more cells per group. Significance denoted by **P* ≤0.05, ***P*≤0.01, ****P*≤0.001 and *****P*≤0.0001, by two way ANOVA with Tukey’s multiple correction test (A), ordinary one way ANOVA with Tukey’s multiple correction test (C to F). **OCR-** oxygen consumption rate, **DMOG**- dimethyloxaloylglycine, **DMOG+AV**- DMOG+10µg/ml. *Adhatoda Vasica* extract, **DMOG+Dex**- DMOG+10nM of Dexamethasone, **DMOG+Dex+AV**-DMOG+10nM of Dexamethasone+10µg/ml. *Adhatoda Vasica* extract, **DMOG 8hrs**- DMOG treatment for 8 hours.

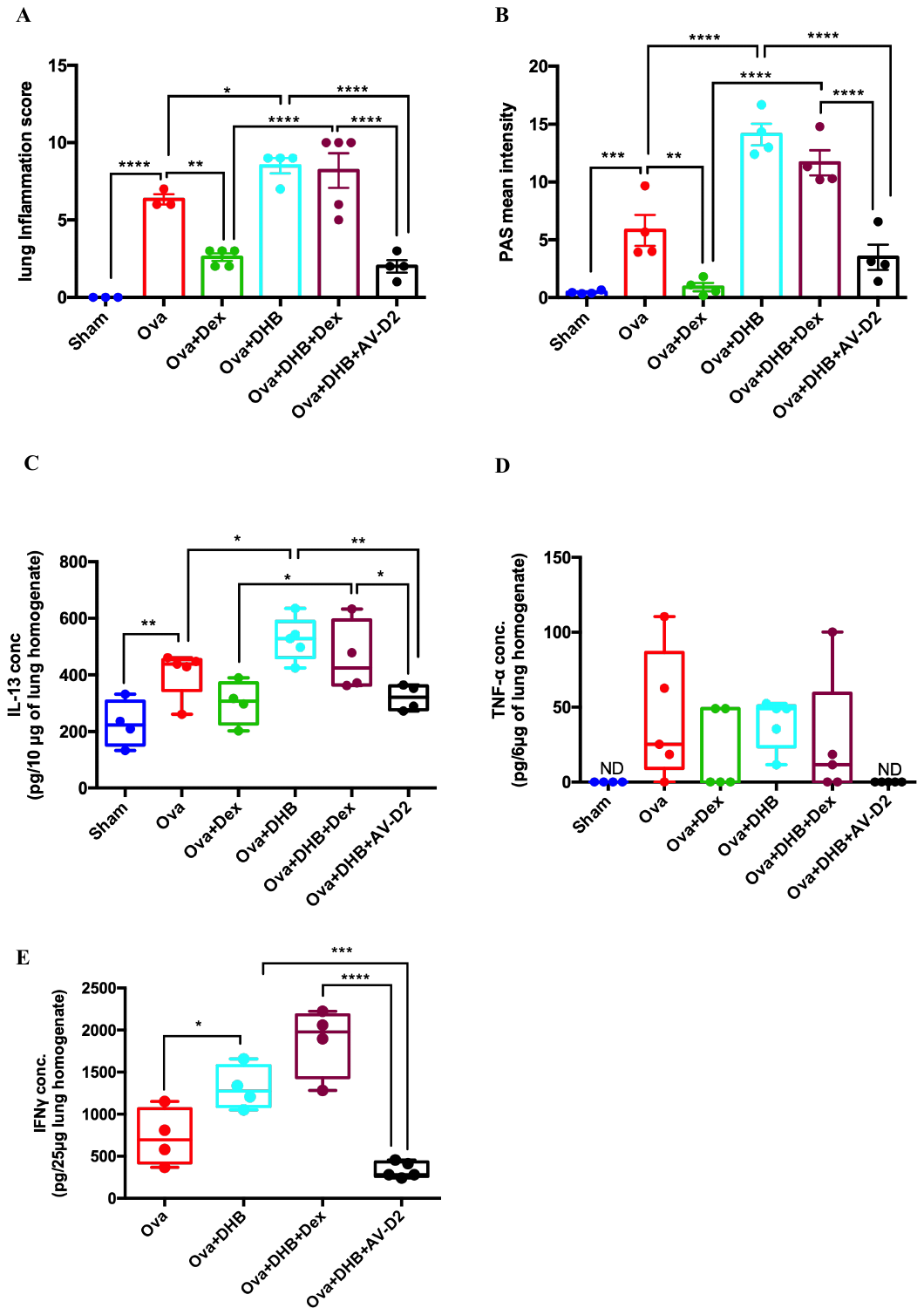

**Figure S4: AV resolves the chemically induced severe corticosteroid insensitive airway inflammation. (A)** Quantification of Peribronchial and Perivascular inflammation of lung tissues stained with H&E by inflammation scoring grade method. **(B**.) Densitometric analysis of mouse lung tissues stained with PAS to measure goblet hyperplasia in severe asthmatic mice using ImageJ **(C to E)** ELISA for IL-13, TNF-α and IFN-γ cytokine levels measured in lung tissue lysate of severe asthmatic mice. Data are shown as mean ±SEM of three to five mice per group and representative of two independent experiment.. Significance denoted by **P* ≤0.05, ***P*≤0.01, ****P*≤0.001 and ****P*≤0.001; by ordinary one way ANOVA. **Ova-** chicken egg albumin, **Sham-** vehicle (PBS), **DHB**- ethyl, 3,4 -dihydroxy benzoic acid (10mg/kg), **Dex-** Dexamethasone (0.75mg/kg), **AV-D2**- *Adhatoda Vasica* extract (130mg/kg).

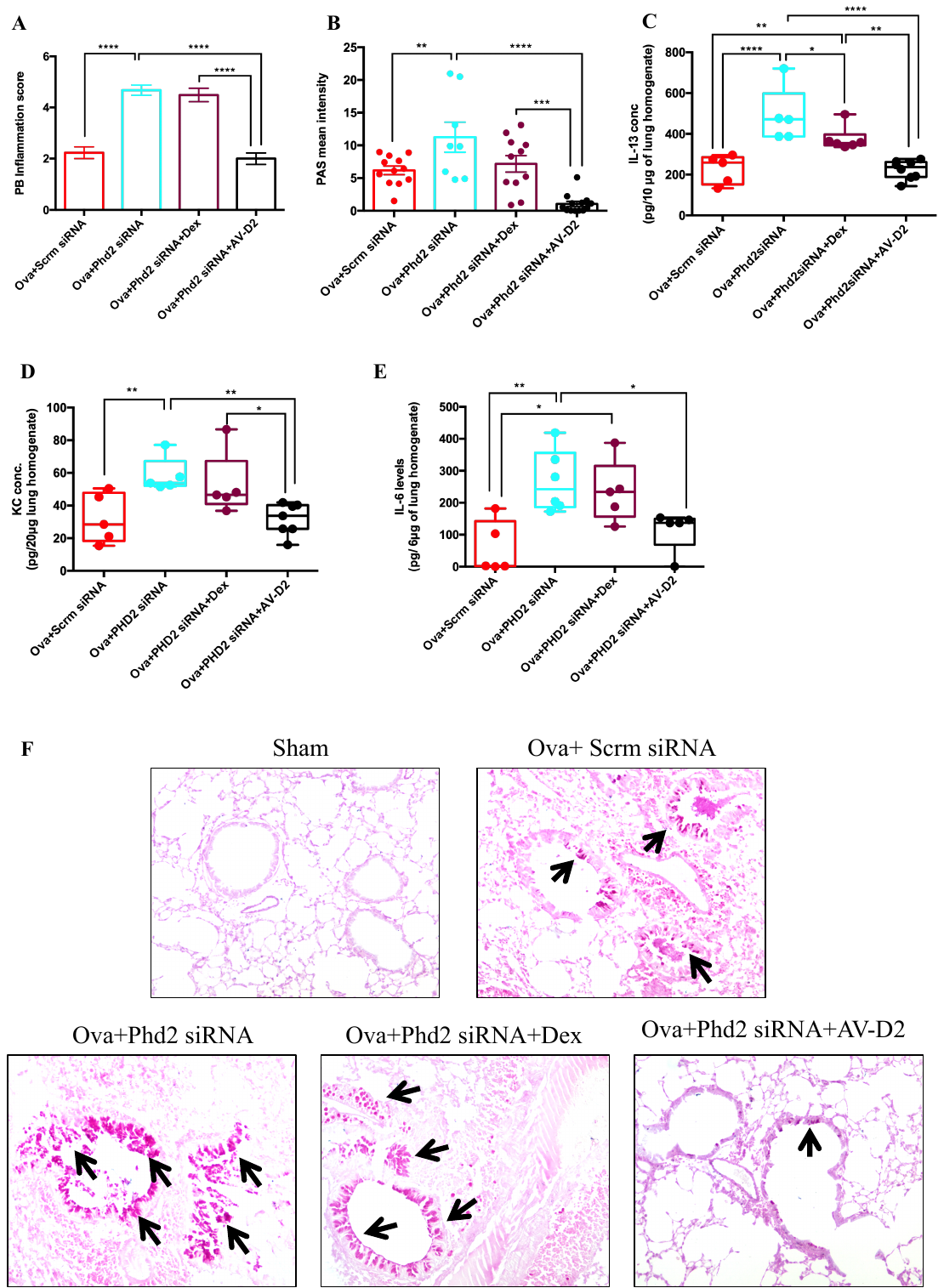

**Figure S5: AV resolves the *PHD2* siRNA induced severe corticosteroid insensitive airway inflammation. (A)** Quantification of Peribronchial inflammation of lung tissues stained with H&E by inflammation scoring grade method. **(B)** Densitometric analysis of mouse lung tissues stained with PAS to mucus metaplasia in severe asthmatic mice using ImageJ. **(C to E)** ELISA analysis of IL-13 (C), KC (D) and IL-6 (E) cytokine levels in lung tissue lysate of severe asthmatic mice. **(F)** Representative photomicrographs of mouse lung tissues stained with PAS to check mucin levels. Arrow indicates positive staining for mucin. Data are shown as mean ±SEM of five to eight mice per group and representative of two independent experiments. Significance indicated by **P* ≤ 0.05, ***P* ≤ 0.01, ****P* ≤ 0.001 and ****P* ≤ 0.001; by ordinary one way ANOVA. **Ova-** chicken egg albumin, **Sham-** vehicle (PBS), **Scrm siRNA-** scrambled siRNA- 90 μg/mice, **PHD2 siRNA-** mouse specific PHD2 siRNA 90μg/mice., **Dex-** Dexamethasone (0.75mg/kg), **AV-D2-***Adhatoda Vasica* extract (130mg/kg)

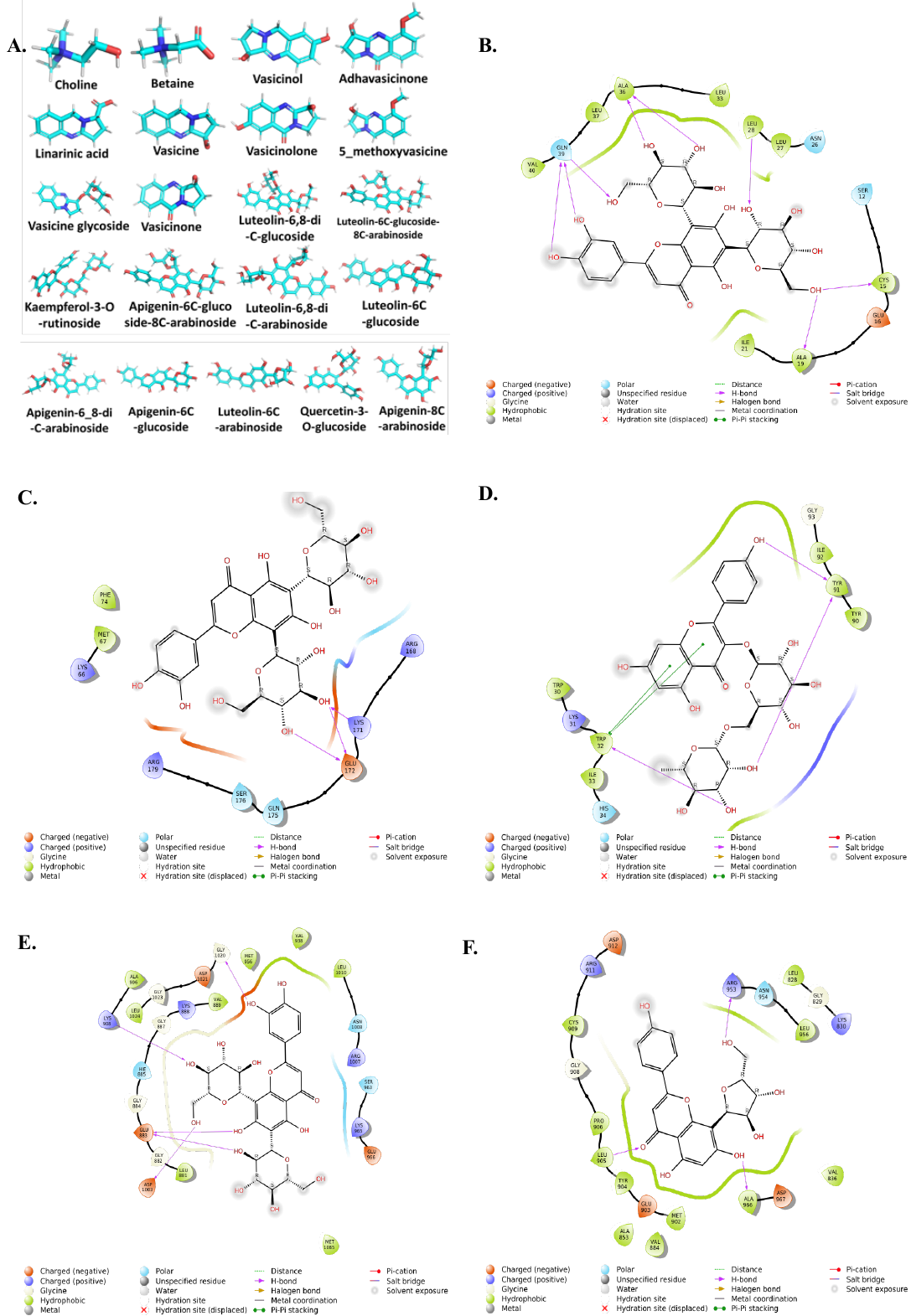

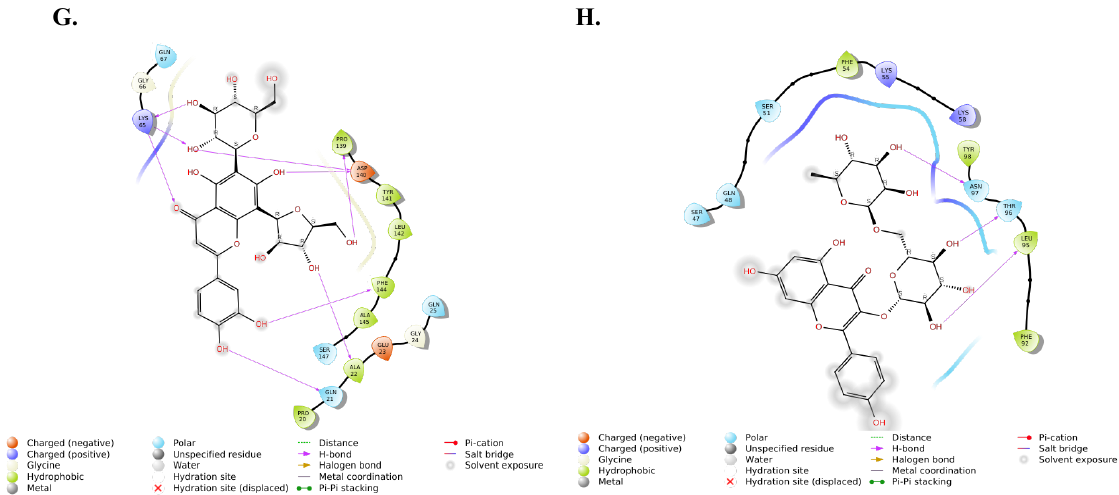

**Figure S6. Interaction of key lead compounds of AV extract with proteins of hypoxia and inflammation pathway.** (**A**) 3-Dimensional structures of constituents of *Adhatoda Vasica*. (**B**) Interaction of Luteolin-6,8-di-C-glucoside with HIF-1α. (**C**) Interaction of Luteolin-6,8-di-C-glucoside with IL-6. (**D**) Interaction of Kaempferol-3-O-rutinoside with TGF-β. (**E**) Interaction of Luteolin-6,8-di-C-glucoside with JAK-1 (**F**) Interaction of Apigenin-8-C-arabinoside with JAK-3. (**G**) Interaction of Luteolin-6C-glucoside-8C-arabinoside with TNF- α. (**H**) Interaction of Kaempferol-3-O-rutinoside with IFN-γ

**Supplementary tables**

**Table S1- Liquid chromatography–electrospray ionization–mass spectroscopy of AV extract in positive-ion mode.** Identification of Quinazoline alkaloids from the water extract of *Adhatoda Vasica,* separated by mobile phase consist of 0.1% formic acid aqueous solution and acetonitrile. The various peaks obtained (fig. S1A) were analyzed at different time intervals under a gradient program using (+)-ESI-MS.

| **S.No.** | **RT  (in min)** | **Molecular Formula** | **[M+H]^+^ *m/z* (calc)** | **[M+H]^+^ *m/z* (exp)** | **Error (Δppm)** | **Identification** |
| --- | --- | --- | --- | --- | --- | --- |
| 1 | 2.4 | C_5_H_14_NO^+^ | 104.1073 | 104.1073 | 0 | Choline |
| 2 | 2.9 | C_5_H_11_NO_2_ | 118.0864 | 118.0864 | 0 | Betaine |
| 3 | 8.0 | C_11_H_12_N_2_O_2_ | 205.0972 | 205.0974 | 0.98 | Vasicinol/ 5-hydroxy vasicine |
| 4 | 8.2 | C_12_H_12_N_2_O_3_ | 233.0921 | 233.0921 | 0.00 | Adhavasicinone |
| 5 | 10.0 | C_11_H_12_N_2_O_2_ | 205.0972 | 205.0974 | 0.98 | Vasicinol/ 5-hydroxy vasicine |
| 6 | 10.4 | C_12_H_12_N_2_O_2_ | 217.0972 | 217.0969 | -1.38 | Linarinic acid |
| 7 | 10.9 | C_11_H_12_N_2_O | 189.1022 | 189.1022 | 0.00 | Vasicine |
| 8 | 13.3 | C_11_H_10_N_2_O_3_ | 219.0764 | 219.0766 | 0.91 | Vasicinolone |
| 9 | 13.7 | C_12_H_14_N_2_O_2_ | 219.1128 | 219.1131 | 1.37 | 5-methoxy vasicine |
| 10 | 15.7 | C_17_H_22_N_2_O_6_ | 351.1551 | 351.1548 | -0.85 | Vasicine glycoside |
| 11 | 15.7 | C_11_H_10_N_2_O_2_ | 203.0815 | 203.0812 | -1.48 | Vasicinone |

**Table S2- Liquid chromatography–electrospray ionization–mass spectroscopy of AV extract in negative-ion mode.** Identification of flavonoids C- and O- glycosides from the water extract of *Adhatoda Vasica*, separated by mobile phase consist of 0.1% formic acid aqueous solution and acetonitrile. The various peaks obtained (fig. S1B) were analyzed at different time intervals under a gradient program using (-)-ESI-MS.

| **S.No.** | **RT  (in min)** | **Molecular Formula** | **[M-H]^-^ m/z (calc)** | **[M-H]^-^ m/z (exp)** | **Error (Δppm)** | **Identification** |
| --- | --- | --- | --- | --- | --- | --- |
| 1 | 14.8, 17.9 | C_27_H_30_O_16_ | 609.1461 | 609.1464 | 0.49 | Luteolin-6,8-di-C-glucoside/ Quercetin-3-O-rutinoside |
| 2 | 15.3, 15.8 | C_26_H_28_O_15_ | 579.1355 | 579.1354 | -0.17 | Luteolin-6-C-glucoside-8-C-arabinoside/ Luteolin-6-C-arabinoside 8-C-glucoside |
| 3 | 15.5, 18.8 | C_27_H_30_O_15_ | 593.1512 | 593.151 | -0.34 | Kaempferol-3-O-rutinoside/ Apigenin-6,8-di-C-glucoside |
| 4 | 15.8, 16.2, 16.5, 16.8, 17.3 | C_26_H_28_O_14_ | 563.1406 | 563.1401 | -0.89 | Apigenin-6-C-glucoside-8-C-arabinoside/ Apigenin-6-C-arabinoside 8-C-glucoside/ Apigenin-6-C-arabinoside 7-O-glucoside |
| 5 | 16.6 | C_25_H_26_O_14_ | 549.125 | 549.1253 | 0.55 | Luteolin-6,8-di-C-arabinoside |
| 6 | 16.7 | C_21_H_20_O_11_ | 447.0933 | 447.0928 | -1.12 | Luteolin-8-C-glucoside/ Luteolin-6-C-glucoside/ Kaempferol-3-O-glucoside |
| 7 | 16.8, 17.2, 17.5, 18.1, 18.5 | C_25_H_26_O_13_ | 533.1301 | 533.1303 | 0.38 | Apigenin-6,8-di-C-arabinoside |
| 8 | 18.1, 18.3 | C_21_H_20_O_10_ | 431.0984 | 431.0983 | -0.23 | Apigenin-6-C-glucoside/ Apigenin-8-C-glucoside |
| 9 | 18.6, 19.3 | C_20_H_18_O_10_ | 417.0827 | 417.0831 | 0.96 | Luteolin-8-C-arabinoside/ Luteolin-6-C-arabinoside |
| 10 | 18.7 | C_21_H_20_O_12_ | 463.0882 | 463.0881 | -0.22 | Quercetin-3-O-glucoside |
| 11 | 19.7, 20.1 | C_20_H_18_O_9_ | 401.0878 | 401.0873 | -1.25 | Apigenin-8-C-arabinoside/ Apigenin-6-C-arabinoside |

**Table S3. AV attenuates increased Th2 cytokines levels.** Levels of IL-4, 5 and 13 levels in lung homogenate measured by ELISA in acute asthmatic mouse. Data are shown as mean, ± SEM. *P ≤0.05, **P≤0.01 and ***P≤0.001 compared Ova mice group (n = 4-7 per group).

| **Mice Group** | **IL-4 levels** | **IL-5 levels** | **IL-13 levels** |
| --- | --- | --- | --- |
| SHAM | 72.75(±13.56)* | 115.08(±34.05)* | 147.04(±29.73)* |
| OVA | 124.02(±11.45) | 220.33(±23.64) | 275.89(±31.69) |
| OVA+DEX | 70.04(±9.77)****** | 108.73(±16.62)** | 124.60(±22.12)** |
| OVA+D0 | 48.26(±9.74)** | 123.15(±24.48)* | 114.79(±20.50)** |
| OVA+D1 | 43.64(±10.69)** | 103.12(±27.12)* | 87.37(±18.69)** |
| OVA+D2 | 64.65(±3.89)** | 102.37(±13.25)** | 89.56(±5.98)** |
| OVA+D3 | 74.76(±13.19)** | 128.54(±14.35)* | 104.20(±14.07)** |
| OVA+D4 | 63.29(±7.35)** | 128.54(±19.09)* | 73.93(±21.42)*** |

**Table S4.** Binding affinity of compounds of *Adhatoda Vasica* extract with proteins of hypoxia and inflammation pathways in kcal/mol using Schrodinger XP Glide package (binding energies of AV’s component with tested proteins more than -6 kcal/mol. indicated by red colour)

| Compounds | HIF-1α | IFN-γ | IL-6 | TGF-β | JAK-1 | JAK-3 | TNF-α |
| --- | --- | --- | --- | --- | --- | --- | --- |
| Choline | -2.21 | -2.35 | -1.93 | -3.3 | -2.47 | -3.448 | -1.87 |
| Betaine | -2.35 | -3.14 | -2.74 | -2.29 | -3 | -3.49 | -2.84 |
| Vasicinol | -1.72 | -3.77 | -2.35 | -3.06 | -6.36 | -7.52 | -2.95 |
| Adhavasicinone | -2.09 | -3.91 | -2.59 | -2.26 | -5.5 | -6.03 | -1.23 |
| Linarinic_acid | -2.04 | -4.35 | -2.23 | -5 | -5.19 | -6.02 | -2.48 |
| Vasicine | -1.9 | -3.92 | -2.15 | -3.67 | -5.53 | -5.41 | -2.22 |
| Vasicinolone | -2.3 | -3.79 | -2.93 | -2.74 | -6.3 | -6.91 | -3.24 |
| 5_methoxyvasicine | -1.95 | -3.66 | -2.05 | -2.04 | -5.3 | -5.81 | -1.23 |
| Vasicine_glycoside | -3.59 | -6.11 | -3.73 | -4.03 | -8.22 | -5.97 | -3.73 |
| Vasicinone | -2.19 | -4.55 | -2.33 | -3.21 | -6.58 | -4.44 | -2.6 |
| Luteolin_6_8_di_  C_glucoside | -9.31 | -7.08 | -6.94 | NA | -12.08 | -10.55 | -8.36 |
| Luteolin_6C_glucoside  _8C_arabinoside | -7.93 | -6.63 | -5.74 | -4.22 | -10.25 | -9.61 | -9.41 |
| Kaempferol_3_O_rutinoside | -7.92 | -7.16 | -5.87 | -7.45 | -7.83 | -6.73 | -5.63 |
| Apigenin_6C_glucoside_  8C_arabinoside | -7.55 | -4.31 | -5.12 | -3.71 | -9.09 | -8.08 | -9.18 |
| Luteolin_6_8_di_  C_arabinoside | -7.63 | -6.1 | -6.64 | -3.94 | -10.21 | -8.86 | -8.55 |
| Luteolin_6C_glucoside | -5.95 | -4.98 | -5.21 | -4.06 | -11.11 | -9.85 | -6.88 |
| Apigenin_6_8_di_  C_arabinoside | -6.45 | -6.55 | -5.68 | -3.24 | -7.78 | -8.33 | -8.19 |
| Apigenin_6C_glucoside | -3.03 | -5.9 | -4.46 | -5.92 | -9.21 | -8.01 | -5.85 |
| Luteolin_6_C_arabinoside | -5.1 | -6.57 | -5.33 | -7.33 | -8.9 | -10.04 | -6.43 |
| Quercetin_3_O_glucoside | -6.27 | -7.11 | -5.89 | -4.77 | -6.75 | -9.21 | -7.7 |
| Apigenin_8_C_arabinoside | -4.8 | -4.94 | -5.02 | -4.5 | -10.3 | -11.28 | -6.24 |

**Table S5.** Interaction of active compounds of *Adhatoda Vasica* extract with proteins of Hypoxia and inflammation pathways.

| **S.**  **No** | **Proteins** | **Compounds** | **Interacting Residues** | **No. of**  **H- bonds** | **Docking Score in kcal/mol** |
| --- | --- | --- | --- | --- | --- |
| 1 | HIF-1α | Luteolin-6,8-di-C-glucoside | Cys 15, Ala 19, Leu 28, Leu 37, Gln 39 | 8 | -9.31 |
| 2 | IFN-γ | Kaempferol-3-O-rutinoside | Leu 95, Thr 96, Asn 97 | 3 | -7.16 |
| 2 | IL-6 | Luteolin-6,8-di-C-glucoside | Lys 171, Glu 172 | 3 | -6.94 |
| 3 | TGF-β | Kaempferol-3-O-rutinoside | Trp 32, Tyr 91 | 3 | -7.45 |
| 4 | JAK-1 | Luteolin-6,8-di-C-glucoside | Glu 883, Lys 908, Asp 1003,  Gly 1020 | 5 | -12.08 |
| 5 | JAK-3 | Apigenin-8-C-arabinoside | Leu 905, Arg 953, Ala 966 | 3 | -11.28 |
| 6 | TNF-α | Luteolin-6C-glucoside-8C-arabinoside | Gln 21,Ala 22, Lys 65, Pro 139, Asp 140, Phe 144 | 9 | -9.41 |
